# A continuum of information-based temporal stability measures and their decomposition across hierarchical levels

**DOI:** 10.1101/2025.08.20.671203

**Authors:** Anne Chao, Robert K. Colwell, Johnny Shia, Simon Thorn, Ming-Yun Yang, Oliver Mitesser, Yu-Ting Huang, Mareike Kortmann, Akira S. Mori, Benjamin M. Delory, Andreas Fichtner, Yuanyuan Huang, Christiane Roscher, Bernhard Schmid, Nico Eisenhauer, Jörg Müller

## Abstract

When biodiversity–stability relationships are assessed from temporal patterns of species (or species assemblages) biomass or other key variables, ecological stability is commonly quantified as the inverse of the coefficient of variation (*CV*) or its square. However, just as biodiversity cannot be fully characterized by a single value, the complexity of temporal stability/invariability cannot be completely captured by one metric, especially since *CV* is disproportionally sensitive to large data values. Ecologists now recognize that species diversity can be fully characterized by a continuum of Hill-number-based measures parametrized by a diversity order *q*≥0, which determines the sensitivity of the measure to species abundance. Building on the intuitive concept that temporal stability can be quantified by the closeness between the data vector and the ideally maximally stable vector, we propose a continuum of information-based measures of temporal stability/invariability parameterized by an order *q*>0. This continuous parameter *q* determines the sensitivity of the measure to the magnitude of biomass or other ecosystem functions. By varying *q*, researchers can differentially weight small, medium, or large values in a time series, thereby disentangling their respective contributions to stability. Our framework unifies and generalizes classical measures: the case *q*=1 links to Shannon entropy, reflecting MacArthur’s 1955 stability concept; *q*=2 connects to the conventional *CV*-based measure. Unlike these traditional metrics, our approach explicitly accounts for the number of data values (i.e., time points), adjusting for time-series-length effects to enable fair and meaningful comparisons across datasets of varying lengths. We extend the framework to hierarchical structures by developing additive and multiplicative decompositions of stability in a metacommunity (or metapopulation) into alpha and beta components. The beta component can be further used to obtain measures that quantify (a)synchrony among communities (or populations). For *q*=2, the resulting (a)synchrony measure provides a mathematically rigorous, *CV*-based metric. The proposed measures are illustrated using 22-year biomass time series data from the Jena Experiment in Germany. We developed the R package iSTAY (information-based stability measures) and online tools for computation and visualization. Our measures are adaptable to other functions beyond biomass and are applicable in both temporal and spatial contexts.

**Open Research Statement:** The original biomass data of the Jena Experiment, covering the years 2003 to 2023, can be retrieved from https://jexis.idiv.de/ddm/Data/ShowData/624, while the 2024 data are available at https://jexis.idiv.de/ddm/Data/ShowData/695. These datasets, along with the R code used in this study, are currently accessible on Github at https://github.com/AnneChao/MS_iSTAY for review purposes and will be archived on Zenodo upon journal acceptance.

## 1. Introduction

Biodiversity−stability relationships have been widely discussed in the literature and have fueled intense debates (McCann, 2000; Loreau et al. 2001; Tilman et al. 2006). Both diversity and ecological stability are multifaceted concepts that can be applied to different levels of organization and across various spatial scales (Mayor et al. 2024; Liang et al. 2025). Ecological stability itself encompasses many concepts (Isbell et al., 2015; Donohue et al., 2016; Loreau, 2022) and multiple mechanisms have been identified that drive biodiversity effects on ecosystem stability (Schnabel et al. 2021; Eisenhauer et al., 2024). One commonly used concept of stability focuses on temporal invariability, i.e., the ability of a system to maintain a consistent state or a constant level over time, typically based on biomass, productivity, or pertinent ecosystem functions/variables (Tilman and Downing, 1994; Isbell et al., 2015). For simplicity in presentation, we use biomass as a representative of a general non-negative variable. Throughout, ecological stability (or its complement, instability) simply refers to invariability (or its complement, variability).

A conventional measure of temporal variability is the coefficient of variation (*CV*) or *CV*^2^, where *CV*^2^ is the ratio of the variance to the squared mean of a biomass series. In the literature, temporal stability is conventionally measured as the inverse of *CV* (or the inverse of *CV*^2^); see, e.g., Pimm (1984), Tilman (1996), Tilman *et al*. (2006), Wagg et al. (2022). The same measure is also used to quantify spatial stability and variability (e.g., Weigelt et al., 2008; Eisenhauer et al., 2011; Wang et al., 2019; Gottschall et al., 2022; Wisnoski et al., 2023).

Just as species diversity cannot be fully characterized by a single number, the complexity of temporal stability cannot be completely captured by a single *CV* value. In particular, using biomass as an example, *CV* is primarily sensitive to large biomass values, implying that it quantifies the temporal variability of large biomass over time. Ecologists now recognize that species diversity can be fully characterized by a continuum of Hill-number-based measures, parametrized by a diversity order *q* ≥ 0, which determines the sensitivity of the measure to species abundance (Chao and Colwell, 2022). Hill numbers encompass the three most widely used measures of species diversity: species richness (*q* = 0), Shannon diversity (*q* = 1, the exponential of Shannon entropy), and Simpson diversity (*q* = 2, the reciprocal of the Simpson concentration index). These three measures are primarily driven by rare (*q* = 0), abundant (*q* = 1) and highly abundant (*q* = 2) species, respectively. This continuum of diversity measures captures all information contained in species relative abundance distributions, in the sense that knowing one set completely determines the other (Chao et al. 2014, Appendix I). Consequently, any change in relative abundance of species will be reflected in the diversity spectrum. In this paper, we aim to develop an analogous continuum of stability measures that encapsulates the complete information contained in a biomass time series.

In Section 2.1, we develop the proposed continuum of stability measures for a single system, parameterized by an order *q* > 0. (Here *q* = 0 is not meaningful, because the stability of *q* = 0 disregards the magnitude of any biomass value.) Intuitively, our approach is to quantify temporal variability as a normalized, information-based distance/divergence between the relative biomass data vector and the maximally stable vector. The corresponding stability is measured by the normalized extent of closeness between these two vectors. The curve/profile that depicts stability with respect to the order *q* > 0 completely characterizes the stability of a system and encompasses all information contained in the biomass time series. Numerical examples are provided in the Applications Section to demonstrate stability profiles. The special cases of *q* = 1 and *q* = 2 are treated in Section 2.2, and each can be linked to conventional stability measures proposed in the literature (see the next three paragraphs).

*Shannon entropy* (or simply *entropy*), originally developed by Claude Shannon in 1948 in information science, was introduced to ecology a decade later by Ramón Margalef as a diversity index (Margalef, 1958). Based on species-abundance or -frequency data, entropy in the ecological diversity context quantifies the uncertainty in the species identity of a randomly chosen individual from a community. Entropy (in units of information) and its exponential (in units of “species equivalents”, MacArthur, 1965) have been applied in many disciplines and played an essential role in the quantification of biodiversity; see Chao and Colwell (2022) and Gotelli et al. (2024) for two recent reviews on the evolution and progress of measuring biodiversity.

As for the quantification of ecological stability, MacArthur in his pioneering work applied Shannon entropy to measure community stability based on pathways for energy flow in a trophic web (MacArthur, 1955). Surprisingly, in contrast to its many applications to biodiversity, Shannon entropy has scarcely been adopted by ecologists to assess stability/invariability of communities or ecosystems; but see Hairston *et al*. (1968) and Colwell (1974).

In the special case of *q* = 1, our measure can be connected to Shannon entropy, whereas the special case of *q* = 2 can be linked to the conventional *CV*^2^. Our framework thus integrates the conventional *CV* measure and MacArthur’s entropy-based modified measure into a single unified family. However, unlike Shannon entropy and the conventional *CV* measure, which do not consider any time-series-length effect, our measures incorporate time-series-length adjustments to address the effect that “more time means more variation” (Lawton, 1988; Pimm and Redfearn, 1988). This adjustment is essential to separate intrinsic ecological responses from confounding effects of time series length. We also show in Section 2.3, using numerical examples, why incorporating time-series-length effects is important.

Thibaut and Connolly (2013) and Wang and Loreau (2014) made important advances by decomposing the *CV* in a metapopulation or metacommunity into within-species (or within-community) variability and among-species (or among-community) asynchronous dynamics. The decomposition framework developed by Wang and Loreau (2014) yielded a synchrony measure that aligns with a widely-used measure proposed by Loreau and de Mazancourt (2008). Subsequently, Gross et al. (2014), Blüthgen et al. (2016), and Lepš et al. (2018) proposed modified versions of this popular synchrony measure.

Analogous to diversity decomposition, we show in Section 3 that the proposed stability measure of any order *q* > 0 in a metacommunity (or metapopulation) can be partitioned additively or multiplicatively into alpha and beta components. These components represent the within-community (or within-population) component and the among-community (or among-population) component. The beta component can further be used to obtain measures to quantify synchrony among communities (or populations). We show that both additive and multiplicative decompositions lead to identical synchrony measures. For *q* = 2, the resulting synchrony measure—which incorporates the effect of number of time points—provides a mathematically rigorous, *CV*-based metric. We also compare our synchrony measures with the one proposed by Loreau and de Mazancourt (2008).

In Section 4, the proposed measures are illustrated using the 22-year biomass time series data collected from the Jena Experiment in Germany (Roscher et al., 2004; Weisser et al., 2017); the biodiversity–stability and biodiversity–synchrony relationships are also examined. We developed the accompanying R package, “iSTAY” (information-based stability measures), and online software to facilitate all computations and graphics. The patterns observed from our measures provide new insights into the mechanisms underlying biodiversity–stability and biodiversity–synchrony relationships in grassland ecosystems.

## 2. A Continuum of Stability Measures in a Single System

### 2.1. Stability measure of a general order *q* > 0

Consider a single level of ecological organization, such as a population, community, or ecosystem. Let {*z*(*t*); *t* = 1, 2,…,*T*} denote a time series of biomass, productivity, or another variable at time point *t*, *t* = 1, 2, …, *T*, where *T* represents the number of time points (i.e., the number of data values), also referred to as the *time series length*. Throughout this article, a “+” sign in place of an index indicates summation over all possible values of that index. For notational simplicity, when the summation is over all time points, *z*(+) is simply denoted as *z*.

For example, the total biomass is represented as 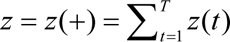 using this simplified notation. This convention extends naturally to cases involving multiple indices in later sections. Let *r*(*t*) = *z*(*t*) / *z* denote the relative biomass at time point *t*, and let ***r*** = {*r*(*t*); *t* = 1, 2,…,*T*} denote the corresponding relative biomass vector. As with the conventional *CV* or its inverse, our variability and stability measures depend only on the values of relative biomass (or another key variable). For simplicity, in this presentation, we focus on the relative biomass time series, recognizing that our stability and synchrony measures apply equally to any other time series of non-negative numbers.

The relative biomass time series is maximally stable (and minimally variable) if and only if biomass remains constant over time, i.e., total biomass is evenly distributed across the *T* time points, with relative biomass values of 1/*T*. Let the mean vector ***r*** = {(1/*T*), …, (1/*T*)} represent the maximally stable case, in which all relative biomass values are equal to 1/*T*. In contrast, the maximally unstable (i.e., minimally stable) series consists of one super-dominant biomass value (i.e., a value whose relative biomass tends toward unity), while the remaining *T*−1 values are vanishingly small. We denote this relative biomass vector with *T* elements as ***r***^0^ = {1^−^, 0^+^, …, 0^+^} where 0^+^ indicates a vanishingly small biomass value and 1^−^ represents a value close to 1. Notably, these two extreme vectors (***r*** and ***r***^0^) are also the two *T*-component vectors that minimize and maximize the conventional *CV*, respectively.

Our intuitive approach is to quantify temporal variability based on a normalized distance/divergence between the relative biomass data vector and the maximally stable vector. Stability is then measured by the normalized extent of closeness between the two vectors, with a greater closeness indicating higher stability. Therefore, if the relative biomass vector represents species relative abundances, our approach is conceptually similar to the framework used to quantify evenness among species abundances in an assemblage within the context of species diversity of Chao and Ricotta (2019). Adapting their approach to the concept of temporal stability, we first define the *temporal information ^q^I* (***r***) of order *q* > 0 based on a relative biomass time series ***r*** = {*r*(*t*); *t* = 1, 2,…,*T*} as

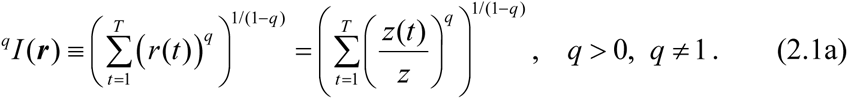

The expression above corresponds to the exponential of Rényi entropy in information science, or Hill numbers in diversity analysis (Hill, 1973). Analogous to the diversity order *q* in Hill numbers, the stability order *q* > 0 in our framework determines the sensitivity of the stability measure to the magnitude of biomass values. The case *q* = 0 is excluded, as it completely disregards biomass magnitudes, rendering the measure meaningless in this context. By varying the parameter *q*, researchers can emphasize small (*q* < 1), medium (*q* = 1), or large (*q* > 1) data values within a time series, allowing them to disentangle the effects of different relative magnitudes on stability dynamics.

There are other forms of information-based measures such as generalized entropy; see the Discussion for justification of our choice. For *q* =1, the limit of the above information reduces to

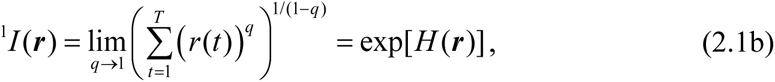

where

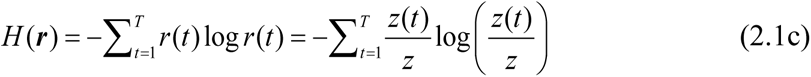

denotes the Shannon entropy based on the relative biomass vector. For *q* = 2, the information ^2^*I* (***r***) reduces to the inverse of the Simpson concentration index when ***r*** = {*r*(*t*); *t* = 1, 2,…,*T*} represents a species relative abundance vector. Let *CV* ^2^ ≡ *CV* ^2^ (***r***) be computed from the relative vector. Then ^2^ *I* (***r***) can be expressed as a decreasing function of *CV*^2^, as shown below:

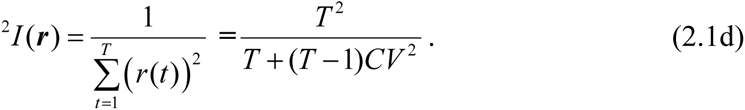

Consider the following *q*th-order (*q* > 0) information-based divergence or distance between the relative biomass vector ***r*** = {*r*(*t*); *t* = 1, 2,…,*T*} and the maximally stable vector ***r̅* = {1(*T*), …,(1/*T***}

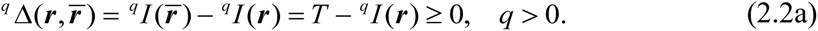

For a general order *q* > 0, the minimum divergence occurs in the special case where ***r̅ = r***, leading to a minimum of zero (min *^q^* Δ(***r***, ***r*** *r̅*) = 0). In contrast, the maximum divergence occurs when one relative biomass value is close to 1 while the other *T*−1 values approach 0, leading to a maximum value of *T*−1 (max *^q^* Δ(***r***, ***r*** *r̅*) = *T* −1). Consequently, we can formulate the normalized divergence as our *variability/instability measure of order q* > 0 as follows:

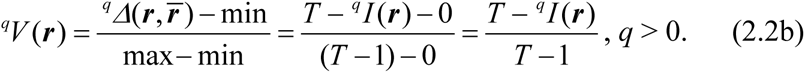

The corresponding *stability/invariability measure of order q* is formulated as the one-complement of *^q^V* (***r***), as follows:

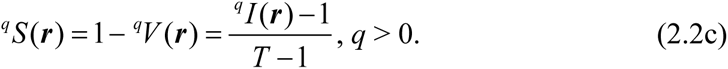

Our stability measure is equivalently based on *^q^I* (***r***) −1 (the numerator in Eq. 2.2c). The denominator *T* − 1 *adjusts for differences in the number of time points* when comparing time-series datasets of varying lengths. The stability measure for any order *q* ranges between 0 and 1. It attains a maximum value of 1 when the time series remains constant and attains a minimum value of 0 when one biomass dominates (i.e., one relative biomass value is close to 1 while all other *T*−1 values are near 0). The special cases of *q* = 1 and 2 are discussed in the next subsection, where we explain the rationale for incorporating the effect of the number of time points into stability analysis.

### 2.2. Special cases: *q* = 1 and 2

In the special case of *q* = 1, the stability measure in Eq. (2.2c) simplifies to

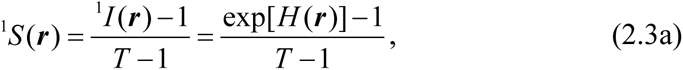

where *H*(***r***) denotes the Shannon entropy as defined in Eq. (2.1c). This measure resembles Heip’s (1974) evenness measure, in which ***r*** = {*r*(*t*); *t* = 1, 2,…,*T*} represents a species relative abundance vector. Shannon entropy is widely used to quantify uncertainty in a system and has applications across various disciplines. However, Shannon entropy alone is insufficient for quantifying stability in a time series because it does not account for the effect of time series length.

Consider all possible relative vectors with *T* components. Shannon entropy attains its maximum value of log*T* when the biomass time series remains constant over *T* time points. Therefore, the range of entropy values depends on the time series length *T*. For instance, when *T* = 3, the maximum entropy is log3 = 1.10 (when biomass is constant over 3 time points); when *T* = 10, the maximum entropy is log10 = 2.30 (when biomass is constant over 10 time points). Consequently, an entropy value of 1.10 represents maximal stability in a series with 3 time points, whereas the same value of 1.10 indicates only moderate stability in a series with 10 time points. This discrepancy creates ambiguity when interpreting entropy magnitudes, and a similar ambiguity arises for any monotonic function of entropy.

Our stability measure with *q* = 1 is based on exp[*H* (***r***)] −1 (i.e., the numerator in Eq. 2.3a), which is a monotonic function of entropy. The denominator *T* − 1 in Eq. (2.3a) *adjusts for differences in the number of time points* when comparing multiple time-series datasets of varying length. This adjustment ensures that the maximally stable time series consistently attains the constant value of 1, while the minimally stable series always attains the constant value of 0, regardless of the time series length. We provide a numerical example in Section 2.3. In the Discussion, we explain why our chosen measure is preferable to the directly adjusted stability formula *H* (***r***) / log *T*.

From Eqs. (2.1d), (2.1b) and (2.2c), it follows that both our stability and variability measures for *q* = 2 are functions of *CV* ^2^ ≡ *CV* ^2^ (***r***) as shown below.

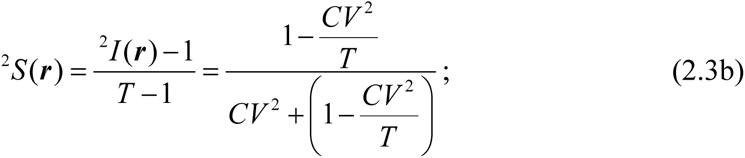

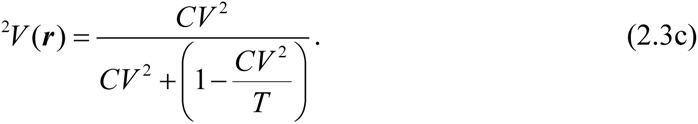

The stability measure derived above is a decreasing function of *CV*^2^, while variability is an increasing function of *CV*^2^. Since our stability measure of *q* = 2 is based on the Simpson index which is disproportionally sensitive to large biomass values, both the conventional *CV*^2^ measure and our *q* = 2 measures primarily quantify the variability or stability of large biomass values within a time series; see Appendix S1 for details.

As noted by Chao and Colwell (2022), using the conventional variability measure (*CV*^2^) to compare datasets with different time series lengths can lead to ambiguity in interpretation. This ambiguity arises because the maximum value of *CV*^2^ for *T* time points is *T* (when one relative biomass value is close to 1 and the remaining *T*−1 values approach 0); see Appendix S1 for a proof. Consequently, the range of *CV*^2^ values depends on the time series length *T*. For instance, when *T* = 2, the maximum value of *CV*^2^ is 2 (one relative biomass value close to 1 and the other close to 0). When *T* = 10, the maximum value is 10 (one relative biomass value close to 1 and the remaining 9 values close to 0). Therefore, a *CV*^2^ value of 2 indicates maximal variability for a series with 2 time points, whereas the same value of 2 for a series with 10 time points indicates low variability. Similar interpretational ambiguity exists for any monotonic function of *CV*^2^.

From Eqs. (2.2c) and (2.1d), our stability measure for *q* = 2 is based on ^2^ *I* (***r***) −1 = *T* ^2^ / [*T* + (*T* −1)*CV* ^2^] −1, which is a monotonic function of *CV*^2^. The denominator *T* – 1 in the stability measure of *q* = 2 (Eq. 2.3b) adjusts for differences in the number of time points. With this adjustment, the maximally stable time series consistently attains the value 1, while the minimally stable series always attains the value 0, irrespective of time series length.

Unlike the conventional *CV*, our measure for *q* = 2 in Eq. (2.3b) accounts for differences in the number of time points. Incorporating the time series length ensures that the same magnitude of our measures consistently quantifies the same degree of variability or stability, even when the length of time series differs. A numerical example is presented in Section 2.3. From the above discussion, a directly adjusted variability formula would be *CV* ^2^ / *T*, with the corresponding stability measure 1− *CV* ^2^ / *T*. In the Discussion, we explain why our chosen stability measure of *q* = 2 (Eq. 2.3b) is preferable to the directly-adjusted formula 1− *CV* ^2^ / *T*.

Unlike the conventional stability measure (1/ *CV*^2^), which becomes infinite when *CV* = 0 (indicating constant biomass over time), our proposed measures of temporal stability and variability are one-complements of each other. As a result, both fall within the range of [0, 1], eliminating the issue of infinity. Specifically, when *CV* = 0, our stability measure attains the maximum of 1, while the variability measure attains the minimum value of 0.

### 2.3 A numerical example to show the importance of incorporating time series length

We present a numerical example to demonstrate intuitively why the number of time points should be considered in the conventional *CV*^2^ measure. Since both *CV*^2^ and our measures are based on relative biomass values, it suffices to use relative vectors in the example. Consider biomass Series A with 2 time points and relative biomass values of 0.999 and 0.001. This series represents almost maximally variable/unstable data for two time points, resulting in *CV*^2^ = 1.992. Next, consider biomass Series B with 10 time points and relative biomass values {3 × 0.3049, 6 × 0.0122, 1 × 0.0121} (i.e., three values of 0.3049, six values of 0.0122, and one value of 0.0121). Although this series deviates significantly from the maximally unstable 10-time-point series, the calculated *CV*^2^ = 1.999 is almost the same as that of Series A. Despite the intuitive expectation of that Series A and B should differ in stability, their inverse *CV*^2^ values are nearly identical, leading to the counterintuitive (and mistaken) conclusion that both series are equally stable.

As discussed in Section 2.2, the above problem arises because the range of *CV*^2^ depends on the number of time points in a time series. According to our proposed measures for *q* = 2, the variability for Series A (Eq. 2.3b) is 0.998 (stability = 0.002), whereas for Series B, the variability is 0.714 (stability = 0.286). This clearly indicates that Series A is less stable than Series B. As shown in Figure S1.1(a) (Appendix S1), for all *q* > 0, our stability measures consistently show that Series A is less stable than Series B, at least to some extent. Similar examples can be found for any other *q* > 0; an example for *q* =1 is provided in Appendix S1.

## 3. Stability Decomposition Across Spatial Scales or Organizational Levels

### 3.1 Alpha and gamma stability

Although Whittaker’s (1960) original concept of species diversity decomposition (alpha, beta, and gamma) pertains to spatial scales or environmental gradients, we adopt his concept of scale as a broad framework for stability partitioning. Specifically, we consider a collection of datasets, each representing a time series of biomass or another key variable. These datasets may originate from sampling units at different levels of ecological organization (e.g., species, guilds, functional groups, habitats, communities or ecosystems) or at various spatial scales (e.g., local, regional, or another spatial unit).

Thibaut and Connolly (2013) and Wang and Loreau (2014) were the first to tackle stability decomposition. Thibaut and Connolly (2013) collected data at the species level (alpha) and aggregated them at the community level (gamma). In contrast, Wang and Loreau (2014) collected data at the local community level (alpha) and aggregated them at the metacommunity level (gamma).

Let *z_k_*(*t*) denote the biomass, productivity or other variable for dataset *k* at time point *t*, where *k* = 1, 2, …, *K* and *t* = 1, 2, …, *T.* In the pooled dataset, let 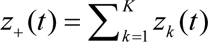 represent the sum of biomass values from all individual datasets at time point *t*. Consider the time series {*z*_+_ (*t*); *t* = 1, 2,…,*T*}. Generalizing the notation in Section 2, we define 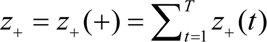 as the total biomass in the pooled dataset. Gamma stability refers to the stability of the pooled dataset. It follows from Eq. (2.2c) that the information-based *gamma stability of order q >* 0 can be formulated using the relative biomass vector in the pooled dataset ***r*** ^(^*^γ^* ^)^ ={*r_γ_* (*t*); *t* = 1, 2,…,*T*}, where *r_γ_* (*t*) = *z*_+_ (*t*) / *z*_+_. Then, gamma stability is

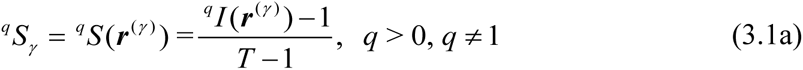

To formulate alpha stability, we first define temporal information and stability in each individual dataset. Let *_q I_* _(***r*** (*k*))_ And *^q^ S* (***r*** ^(*k*)^), *k* = 1, 2, …, *K*, denote, respectively, the temporal information and stability of the *k*th dataset based on the relative biomass vector ***r*** ^(*k*)^ ={*r* (*t*); *t* = 1, 2,…,*T*}, where *r* (*t*) = *z* (*t*) / *z*, and 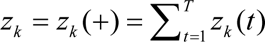 denotes the total biomass in the *k*th dataset. We define the relative-biomass-weight (or simply the *weight* for dataset *k*) as *w_k_* = *z_k_* / *z*_+_, representing the proportion of total biomass contributed by dataset *k* to the pooled dataset. Following the derivations by Routledge (1979) and Chao et al. (2019), we formulate alpha information as a weighted generalized mean of within-dataset information. The information-based *alpha stability of order q >* 0 is then obtained as the following function of weights and the temporal information/stability of individual datasets:

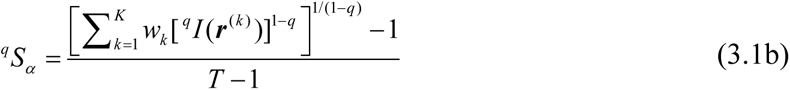

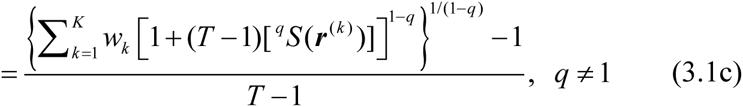

The two special cases of *q* = 1 and *q* = 2 are discussed in Section 3.3.

### 3.2 Beta stability and (a)synchrony

In Appendix S1, we prove that gamma stability is never less than alpha stability for all *q* > 0. Both additive and multiplicative decompositions can be applied, as we later demonstrate that both lead to the same measure of (a)synchrony. For simplicity, we define beta stability as the difference between gamma and alpha stability:

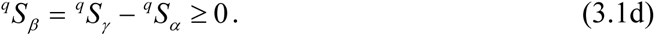

The minimum value of 0 is attained when all samples have identical relative biomass time series, i.e., ***r*** ^(1)^ = ***r*** ^(2)^ = … = ***r*** ^(*K*)^. Given that alpha stability is fixed—that is, given the collection of *K* biomass vectors from individual datasets—the maximum beta stability (denoted as max(*^q^S_β_*) in the following equation) for *q* > 0 is data-dependent and derived as follows (see Appendix S1 for a proof)

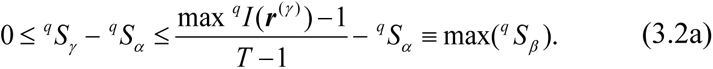

Here max *^q^I* (***r*** ^(^*^γ^* ^)^) = min{*^q^I* (***w*** × ***r***), (1−1 / *K*)*T* + (1 / *K*)[*^q^I* (***w*** × ***r***)]}, where *^q^I* (***w*** × ***r***) denotes the information computed from the two-dimensional *K* × *T* table [*z_k_* (*t*) / *z*_+_] = [*w_k_ r_k_* (*t*)], i.e.,

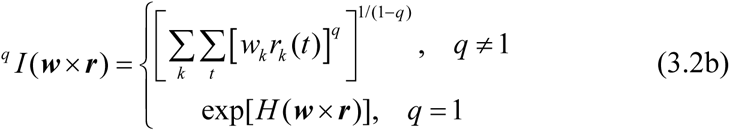

where

*H* 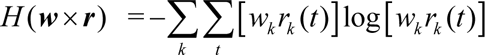 denotes Shannon entropy based on [*w_k_ r_k_* (*t*)]. The upper bound, max(*^q^S_β_*), in Eq. (3.2a) is attained if and only if there exists a dataset λ such that *z***_+_** (*t*) = *z*_<συβ>λ <*t*_(*t*). In other words, at any given time point, the total biomass in the pooled dataset is almost exclusively contributed from a single dataset, while the biomass contributions from other datasets are negligible. Two numerical examples that illustrate how the maximum is obtained are provided in Appendix S2.

Because the range of additive beta stability is data-dependent, comparing beta stability across datasets is not meaningful. To make meaningful comparisons, beta stability values must be normalized to a fixed range of [0, 1] using the following transformation.

The resulting normalized beta stability is our proposed *asynchrony* measure among *K* datasets:

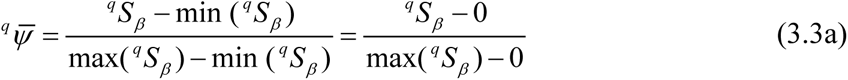

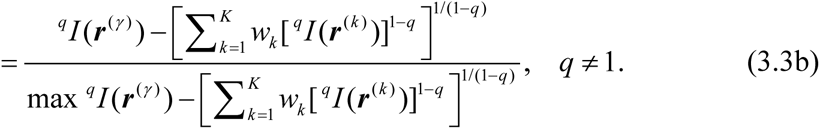

This asynchrony measure is constrained to fall within the range [0, 1]. See Section 3.3 for the special case of *q* = 1. The corresponding *synchrony* among *K* datasets for *q* ≠ 1 is formulated as follows.

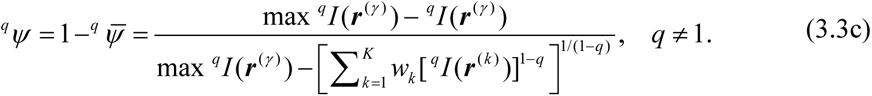

This synchrony measure also falls in the range of [0, 1]. It attains the minimum value of 0 when, at any given time point, the total biomass in the pooled dataset is almost exclusively contributed from a single dataset, while the biomass contributions from other datasets are negligible. It attains the maximum value of 1 when all samples have identical relative biomass time series vectors, i.e., ***r*** ^(1)^ = ***r*** ^(2)^ = … = ***r*** ^(*K*)^. Our synchrony measure thus quantifies *the extent of similarity among relative biomass vectors, while the asynchrony measure quantifies the extent of dissimilarity among relative biomass vectors*.

We can also define multiplicative beta stability as the ratio of gamma to alpha stability, i.e., 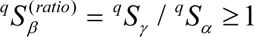. Its minimum is 1, while its maximum is data-dependent. The normalized beta stability (serving as a measure of asynchrony) is then given by

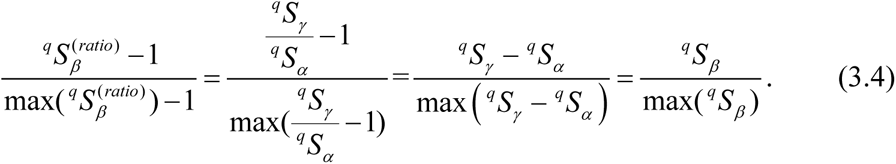

Comparing Eqs. (3.3a) and (3.4), we conclude that both additive and multiplicative decompositions lead to the same normalized formula for (a)synchrony measures.

### 3.3. Special cases: *q* = 1 and 2

Let *H* (***r*** ^(^*^γ^* ^)^) and *H* (***r*** ^(*k*)^) denote, respectively, the Shannon entropy based on the relative biomass vector in the pooled dataset and in the *k*th dataset*, k* = 1, 2, …, *K.* For the special case of *q* = 1, gamma and alpha stability can be expressed in terms of these entropy values as follows:

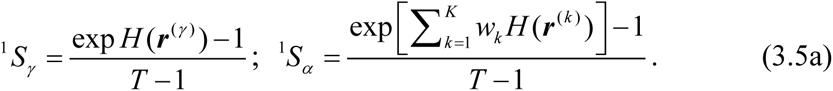

From Eq. (3.3c), the corresponding measure of synchrony among datasets is given by:

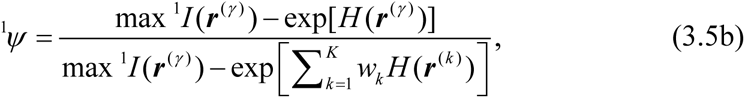

Where max ^1^*I* (***r*** ^(^*^γ^* ^)^) is defined in Eq. (3.2a).

For the special case of *q* = 2, let *CV* ^2^ (***r*** ^(^*^γ^* ^)^) and *CV* ^2^ (***r*** ^(*k*)^) denote, respectively, the *CV*^2^ values based on the relative biomass vector in the pooled dataset and in the *k*th dataset*, k* = 1, 2, …, *K.* From Eqs. (2.3b) and (3.1a), it follows that gamma stability for *q* = 2 can be expressed as the following function of *CV* ^2^ (***r*** ^(^*^γ^* ^)^) :

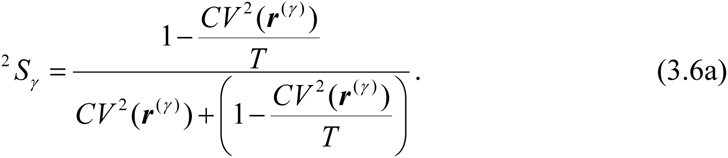

Let ^2^*S* (***r*** ^(*k*)^) denote the stability measure of the *k*th dataset, *k* = 1, 2, …, *K*. For the special case of *q* = 2, Eq. (3.1c) reduces to the following alpha stability formula:

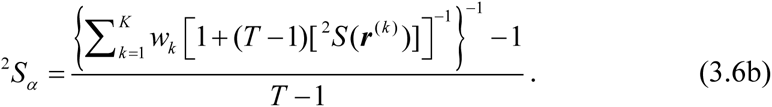

Since ^2^*S* (***r*** ^(*k*)^) is a function of *CV* ^2^ (***r*** ^(*k*)^) (as shown in Eq. 2.3b), alpha stability for *q* = 2 is thus a function of the *CV*^2^ values of individual datasets. The corresponding measure of synchrony among datasets can be obtained by substituting *q* = 2 into Eq. (3.3c). The resulting formula is a complicated function of *CV* ^2^ (***r*** ^(^*^γ^* ^)^), *CV* ^2^ (***r*** ^(*k*)^), *k* = 1, 2,…, *K*, and *CV* ^2^ (***w*** × ***r***) computed from the matrix [*w_k_ r_k_* (*t*)].

In Appendix S3, we compare our synchrony measures for *q* = 1 and 2 with the conventional measure proposed by Loreau and de Mazancourt (2008). The numerical results demonstrate that our synchrony measures offer notable improvements over the conventional measure. See the Discussion section for details.

## 4. Applications

We illustrate the proposed stability measures and their decompositions using biomass time series data collected from the (ongoing) Jena Experiment. For detailed information on the experimental design and dataset, see Roscher et al. (2004), Weisser et al. (2017), and Wagg et al. (2022). The experiment is based on a species pool of 60 plant species belonging to four functional groups: 12 legumes, 16 grasses, 20 tall herbs, and 12 small herbs. Plots were sown with a gradient of 1, 2, 4, 8, or 16 species, originally in 20 × 20 m plots (reduced to 6 × 6 m after 2009), distributed in four blocks.

Our analysis is based on time series data from 2003 to 2024 across 76 plots, with 19, 19, 20, and 18 plots in the four blocks, respectively. See Appendix S4 (Table S4.1) for the distribution of species richness and plots counts across blocks. All feasible combinations of species richness and functional groups were replicated four times with different species compositions, with two exceptions: (1) the 16-species mixtures with one functional group were replicated only twice due to insufficient numbers of species in legumes and smaller herbs; (2) to account for higher variability at low richness levels, there were 14 monoculture plots (two monoculture plots were abandoned in 2009 due to low target species cover), each containing only one functional group. For 2-species mixtures, there are 8 plots with species from a single functional group and another 8 plots with species from two functional groups. See Table 3 of Roscher et al. (2004) for all combinations.

The biomass data from 2003 to 2023 were retrieved from https://jexis.idiv.de/ddm/Data/ShowData/624, while the 2024 data were obtained from https://jexis.idiv.de/ddm/Data/ShowData/695. Biomass was recorded at the species level, except that in 2004 only the pooled biomass of sown species was available. A total of 34 missing biomass data (approximately 0.35% of the dataset) were imputed using the random forest method implemented in the *missForest* package (Stekhoven, 2022). In our analysis, each plot is treated as a community, and the terms “community” and “plot” are used interchangeably (note that each plot had a unique species composition). Community-level biomass was calculated by summing the biomass of all constituent species within each plot from two annual harvests at peak biomass before mowing (June, September). We use 22 years of data (2003−2024) for community-level analysis. For species-level analysis, only 21 years are included, excluding 2004 due to the absence of species-specific biomass records.

In Section 4.1, we illustrate the decomposition theory for partitioning stability within a plot, using two example plots. Section 4.2 presents biodiversity–stability relationships across all 76 plots. Section 4.3 examines temporal effects of species richness on stability and (a)synchrony. In sections 4.1−4.3, time series data are associated with species in a plot (with the plot representing the gamma level), allowing the quantification of species (a)synchrony within plots. In Section 4.4, we examine biodiversity–stability relationships across 20 sets of plots, each treated as a metacommunity composed of plots with equal species richness. In this context, time series are associated with plots within each metacommunity (with the metacommunity representing the gamma level), enabling the quantification of spatial (a)synchrony among plots within metacommunities.

### 4.1 Stability profiles and stability decomposition within a local plot/community

We selected two plots—Plot B1A04 (with 4 species from 4 functional groups) and Plot B4A14 (with 2 species of the same functional group)—to illustrate stability profiles. We also demonstrate the proposed stability decomposition approach within each plot, linking species- and community-level stability, and assess species synchrony within each plot. Biomass time series data for individual species in these two plots are provided in Appendix S4 (Tables S4.2 and S4.3). Under the broad partitioning framework described in Section 3.1, gamma stability corresponds to the plot/community level, alpha stability to the species level, and synchrony to species synchrony within a plot.

Figure 1 displays the gamma, alpha and synchrony profiles for *q* between 0 and 2 within each plot. Figure 1(a) shows the community-level (gamma) stability profiles for the two selected plots. Figure 1(b) presents the species-level stability profiles (dashed lines) for the four species in Plot B1A04 and two species in Plot B4A14; the alpha stability profile (solid line) of each plot is derived as a function of stability profiles of its constituent species (Eq. 3.1c). Notably, the two gamma profiles cross at *q* = 1.4, indicating that different values of *q* (e.g., *q* = 1 *vs. q* = 2) may yield different rankings of stability between plots. In contrast, the two alpha profiles do not cross.

**Figure 1.**
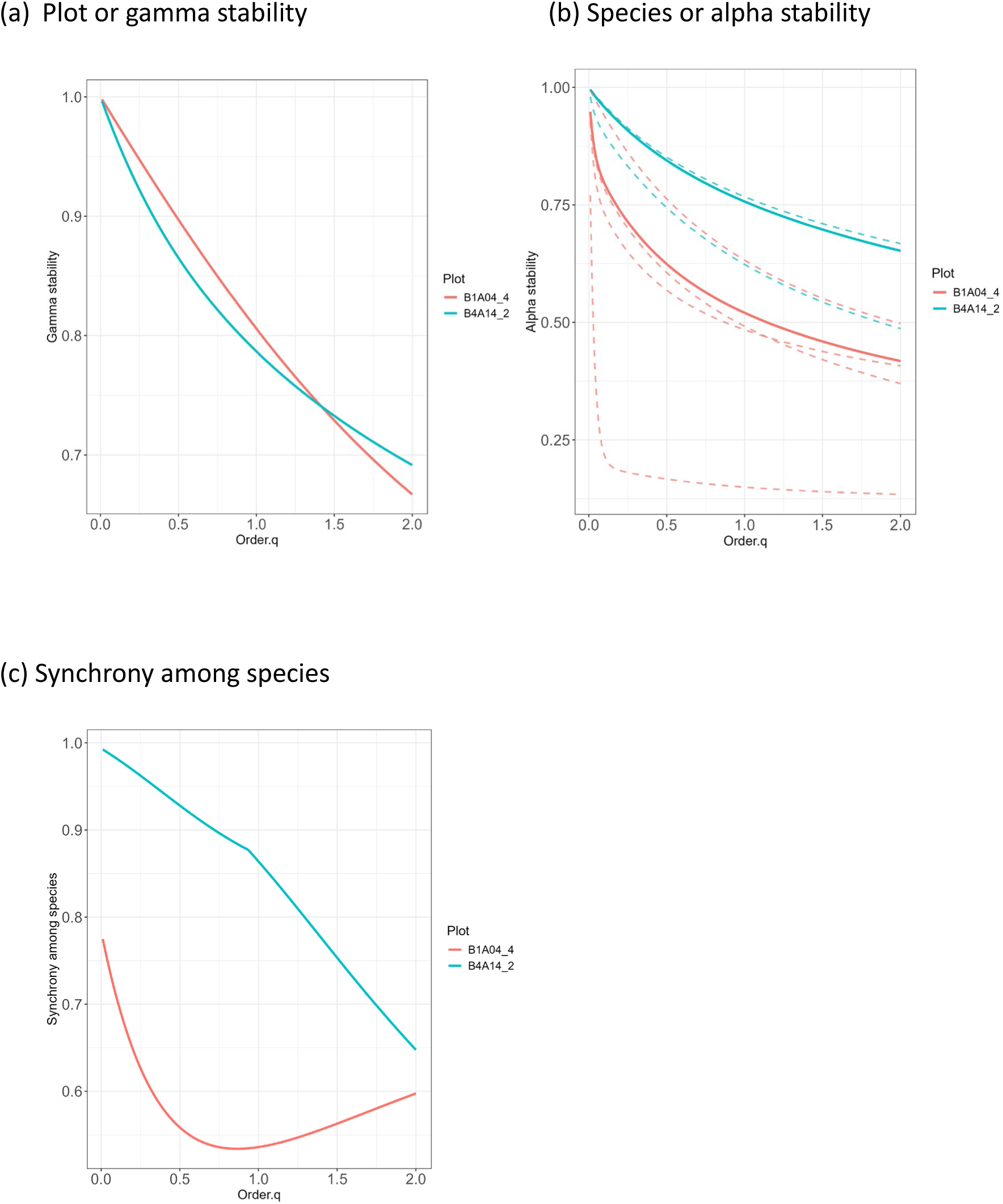
Stability profiles at (a) the gamma/plot level, (b) the alpha/species level, and (c) species synchrony within a plot, for *q* between 0 and 2, based on the data from Plot B1A04 (4 species) and Plot B4A14 (2 species); see Appendix S4 for time series of constituent species from 2003 to 2024. In (a), the two gamma profiles represent the stability of the two plots. In (b), alpha stability is shown as a function of the stability profiles of individual species, with red dashed lines for species in Plot B1A04 and blue dashed lines for species in Plot B4A14.

As discussed in Section 3.2, directly comparing beta stability profiles is not meaningful because their ranges are data-dependent. A valid comparison should instead be based on synchrony measures, i.e., one-complement of normalized beta stability. Figure 1(c) reveals that species synchrony in the 2-species Plot B4A14 is consistently higher than in the 4-species Plot B1A04 for all orders of *q* between 0 and 2.

One advantage of using a continuum of measures is that it allows us to disentangle the effects of large, medium and small biomass values on stability and species synchrony. A consistent pattern observed in the species- and community-level stability profiles is their decline with increasing order *q* (Fig. 1a and 1b). These declining trends suggest that, within a population or community, large biomass values (emphasized at *q* = 2) in a time series tend to be less stable than medium or small values (emphasized at lower *q*). The two synchrony profiles exhibit distinct patterns. In the 2-species Plot B4A14, synchrony decreases with increasing *q*. In contrast, the synchrony profile in the 4-species Plot B1A04 exhibits a U-shaped pattern, where synchrony between species is lowest when emphasizing medium biomass values, but the synchrony is higher when focusing on either small or large biomass values.

### 4.2 Biodiversity–stability and biodiversity–synchrony relationships based on 76 plots

We extended the same decomposition procedures applied to Plots B1A04 and B4A14 (Section 4.1) to all 76 plots. For each plot, we computed alpha and gamma stability values, along with among-species synchrony for *q* between 0 and 2. Since it is not feasible to compare 76 stability and synchrony profiles, we specifically extracted values at three representative *q* values: 0.5, 1, and 2. For each *q*, the biodiversity–stability relationships at both the gamma and alpha scales, along with biodiversity–synchrony relationships, are shown in Figure 2. Note that *q* = 0 is necessarily excluded, as it disregards biomass magnitudes.

**Figure 2.**
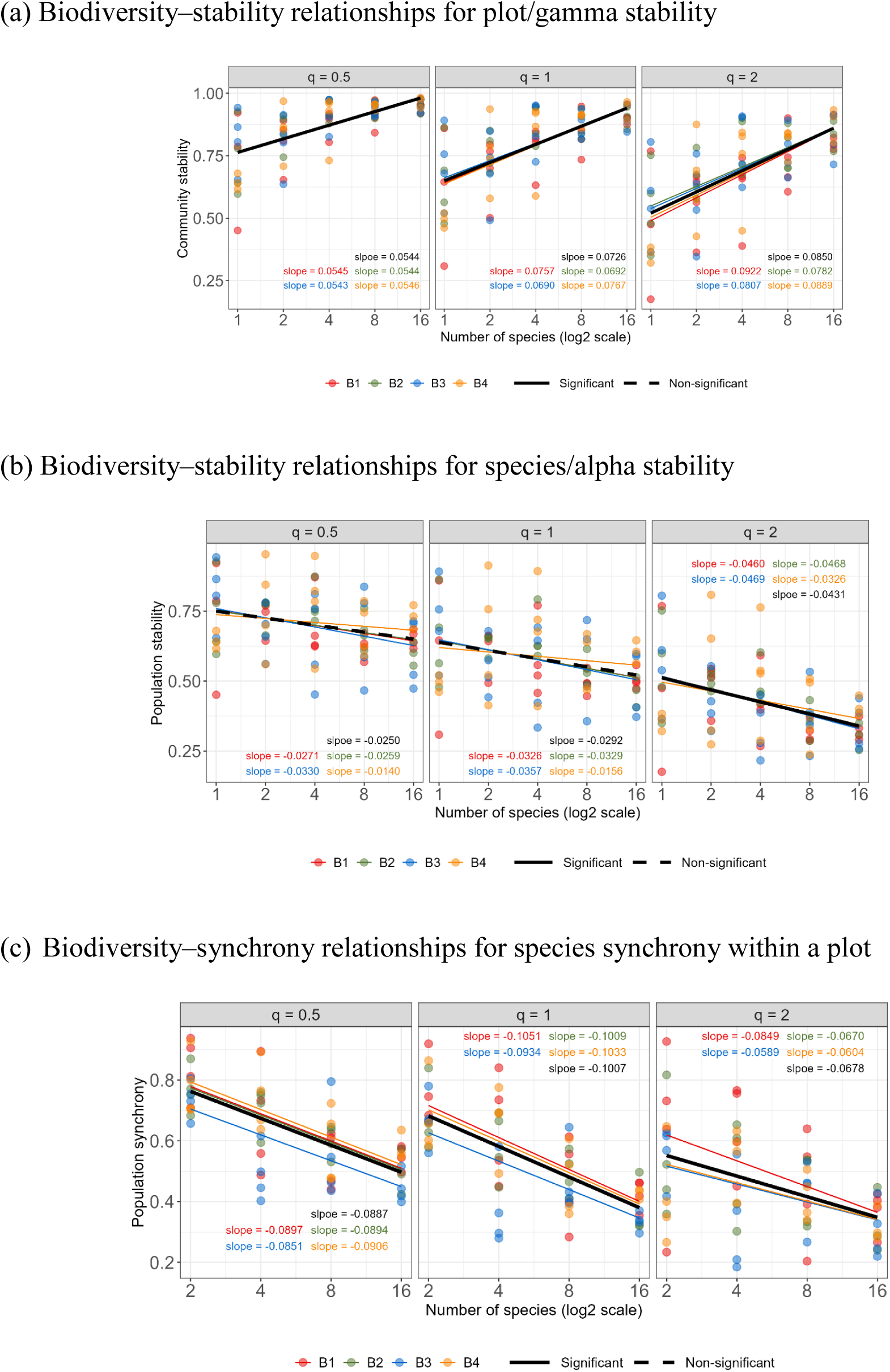
Relationships between the logarithm of species richness and gamma stability (a), alpha stability (b), and species synchrony (c) for orders *q* = 0.5, 1 and 2, based on decomposition within individual plots/communities. In (c), plots with one species were not included, as synchrony is not meaningful for a single time series. Thus, panels (a) and (b) are based on 76 plots, while Panel (c) includes a subset of 62 plots. The relationships between species richness and each measure were modeled using a linear mixed-effects model with random slopes and random intercepts for each block. The figure shows overall fixed-effects slopes (bold black lines) and block-specific relationships (thin lines), as estimated from the same linear mixed-effects model. Because some thin lines overlap with the corresponding bold line, they are not visible. A solid line in the overall fit indicates significance (*P* < 5%), while a dashed line indicates non-significance.

The relationships between biodiversity and gamma stability, alpha stability, and synchrony for each fixed value of *q* were modeled using linear mixed-effects models with random slopes and random intercepts for each block. Figure 2 reveals the overall fixed-effect slopes (bold black lines) along with block-specific relationships (thin lines), as estimated from the same linear mixed-effects model. All fitted results and corresponding significance tests for the overall slopes were obtained using the “lmer” function from the R packages *lme4* (Bates et al., 2015) and *lmerTest* (Kuznetsova et al., 2017).

Figure 2(a) shows that for all values of *q*, gamma stability (community stability based on aggregate biomass) is *positively* related to species richness across the 76 plots. In contrast, Figure 2(b) shows that alpha stability (within-species population stability) is *negatively* related to species richness. The slope is significant at *P* < 5% for *q* = 2 in Figure 2(c). Although this study uses novel stability measures, the observed directions of biodiversity effects on community- and population-level stability are consistent with many previous theoretical and empirical findings. These studies generally suggest that biodiversity enhances community stability but reduce species-level population stability in experimental grasslands (e.g., Roscher et al., 2011, Hautier et al. 2014; Isbell et al., 2015; Xu et al., 2021, Jiang and Xu, 2022, Wagg et al. 2022, Eisenhauer et al., 2024).

Most previous studies on biodiversity–stability relationships have relied on the inverse of the *CV*, which (as we have shown) is primarily sensitive to large biomass values. By employing a continuum of stability measures, our study demonstrates that the opposite direction of biodiversity–stability relationships at community and species levels are also valid when the focus is shifted to intermediate or low biomass values within a time series. Gamma stability exhibits considerably less variability among plots at higher species richness levels, regardless of the value of *q*. A similar pattern is also observed for alpha stability when *q* = 2. Additionally, comparing the fitted linear trends across the three different values of *q* in Figures 2(a) and 2(b), we observe that the magnitude of slope increases with *q*. This pattern signifies that the effect of biodiversity on stability is stronger for larger biomass values —whether in community biomass (Figure 2a) or population biomass (Figure 2b)—than for medium or small biomass.

In our assessment of biodiversity–synchrony relationships (Figure 2c), monoculture plots were excluded because among-species synchrony is not meaningful in plots with only one species. Applying our proposed synchrony measures to species-mixture plots, Figure 2(c) reveals that species in more diverse plots tend to be less synchronized than those in less diverse plots. This *negative* effect is significant for all values of order *q*. In the literature, various measures have been used to quantify different aspects of among-species “synchrony,” leading to contrasting biodiversity–stability relationships and divergent interpretations (e.g., Houlahan et al. 2007, Valencia et al. 2020). In our framework, synchrony (or its complement, asynchrony) quantifies the extent of similarity (or its complement, dissimilarity) among relative biomass time series.

Under our proposed decomposition framework, temporal stability of any order *q* > 0 in a community is determined by within-species population stability (alpha component) and among-species asynchronous dynamics (beta component). In the Jena Experiment, as biodiversity increases, species-level population stability declines (Figure 2b) for all orders of *q*. Therefore, the observed increase in community stability (Figure 2a) in more diverse communities must be driven by a corresponding increase in species-level asynchrony—reflected by the decrease in synchrony shown in Figure 2(c). This pattern confirms that greater biodiversity enhances community stability primarily through increased asynchrony among species (Wagg et al. 2022).

### 4.3 Temporal effects of species richness on stability and synchrony based on 12 consecutive, overlapping 10-year moving window

In Sections 4.1 and 4.2, all measures were calculated using the complete 22-year biomass time series from the Jena Experiment. To investigate the temporal effects of species richness on community stability, species-level population stability, and species synchrony, we further adopted a moving-window approach, following Wagg et al. (2022). Unlike their use of a 5-year moving window, we employed a 10-year moving window to reduce the influence of short-term fluctuations that could obscure underlying patterns. Due to the unavailability of species biomass data in 2004, this year was excluded from the analysis. Consequently, the first time period covers 2003 and 2005–2013, the second period spans 2005–2014, the third period covers 2006–2015, and so on, resulting in a total of 12 overlapping decades.

Within each time period, we first computed the average gamma (community-level) stability and alpha (species-level) stability for orders *q* = 0.5, 1, and 2 across all plots with the same number of species. Since synchrony is not meaningful for a single time series, average species synchrony values were calculated only among plots containing species mixtures. The left panels of Figure 3 illustrate how the average values of gamma stability, alpha stability, and synchrony vary across the 12 moving windows for each value of *q*. In general, for more diverse plots (species richness of 4, 8, or 16), the average gamma stability remains relatively constant over time. In contrast, for less diverse plots (species richness of 1 or 2), alpha and gamma stability decline rapidly through time. Regardless of species richness, average species synchrony shows little variation during the first half of the study period but exhibits a gradual decreasing trend in the later half.

**Figure 3.**
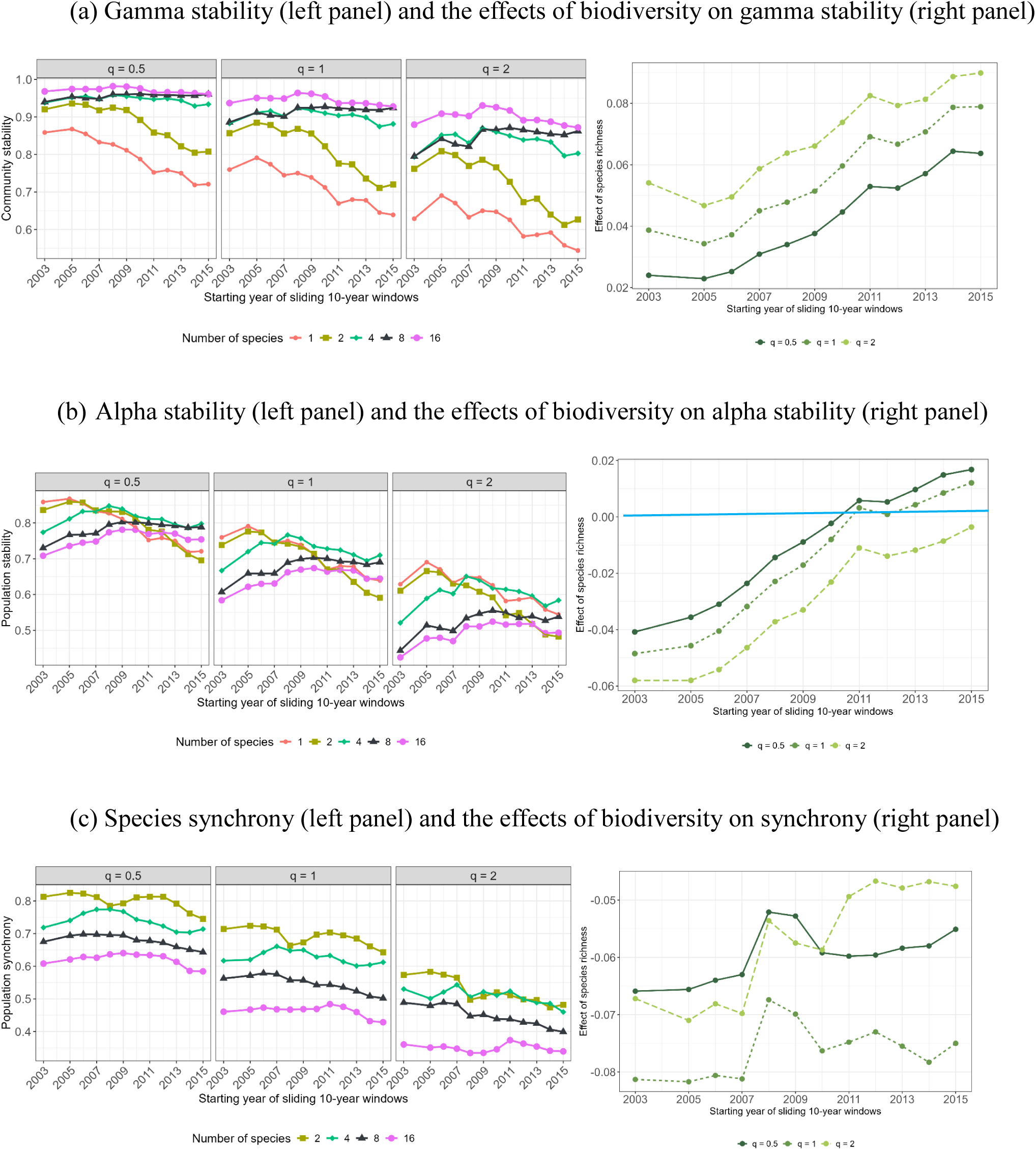
(Left panels) The average (a) gamma (plot-level) stability and (b) alpha (species-level) stability of order *q* = 0.5, 1, and 2 across all plots sharing the same number of species. The average species’ synchrony values (c) were obtained across plots with monoculture plots excluded. (Right panels) The effects of species richness on (a) gamma stability, (b) alpha stability, and (c) species synchrony, represented by the slopes of each biodiversity–stability or biodiversity–synchrony relationship across 12 consecutive 10-year windows. In Panel (a), all effects are positive (note the Y-axis range) and effect increases over time. In panel (b), the magnitude of negative effects diminishes over time and becomes positive in the last five periods, for *q*=0.5 and *q*=1. The blue horizontal line indicates a neutral effect (effect = 0). In Panel (c), all effects are negative, and the magnitude of the negative slope exhibits a decreasing trend.

Within each time period, we separately fitted linear mixed-effects models following the procedures described in Section 4.2 to assess the effects of species richness on gamma stability, alpha stability, and species synchrony. As it is impractical to present fitted lines for all 12 time periods, we show the relationships for three representative periods in Appendix S4. These periods are defined as follows: an early period (2005–2014), a middle period (2010–2019), and a late period (2015–2024). The early and late periods are non-overlapping, while the middle period overlaps with each of the other two by five years. Readers are referred to these three periods (Figure S4.1) to facilitate a clearer understanding of the general conclusions summarized below, based on analyses across all 12 consecutive 10-year periods.

The right panels of Figure 3 illustrate how the slopes of the biodiversity–stability and biodiversity–synchrony relationships vary across 12 consecutive 10-year windows. These slopes represent the effect of species richness on stability and synchrony. Consistent with the patterns shown in Figures 2(a) and 2(c), the right panels of Figures 3(a) and 3(c) demonstrate that species richness has a consistently positive effect on gamma stability and a consistently negative effect on species synchrony, regardless of the time period or the value of *q*. For any fixed value of *q*, comparisons across time periods reveal that the stabilizing effect of species richness on gamma stability strengthens over time, as indicated by increasingly positive slopes—a pattern also reported by Wagg et al. (2022), although their analysis was based on different stability measures and shorter time series. In contrast, Figure 3(c) shows that the magnitude of the negative effect on species synchrony weakens over time, reflected by a decreasing magnitude of the negative slopes; equivalently, the positive effect on species asynchrony becomes weaker over time. Figure 3(b) shows that in the early and middle periods, alpha stability is negatively related to species richness, consistent with the pattern observed in Figure 2(b). However, in the later periods, the magnitude of this negative effect diminishes for *q* = 2 (Wagg et al. 2022); for *q* = 0.5 and 1, the effect even reverses and becomes positive in the last five periods.

These results suggest that the effects of species richness on gamma stability, alpha stability and species synchrony change progressively with time. Consequently, as communities age, the mechanisms underlying community stability shift (Wagg et al. 2022). In the early periods, species asynchrony is the primary driver of gamma stability, as species richness negatively affects alpha stability. Over time, the negative effects of diversity on species stability diminish or vanish, while the positive effect on species asynchrony weakens. (By the later periods, for *q* = 0.5 and 1, both species asynchrony and species stability contribute to enhancing community stability.) Taken together, these dynamical processes result in a strengthening overall effect of species richness on gamma stability over time. Unlike Wagg et al.’s finding that the effect on gamma stability seemed to taper off after the first decade, Figure 3(a) reveals that the stabilizing effect continues to increase over time, regardless of the order of *q*. In addition, the effect increases with the order *q* within any time period, indicating that the stabilizing effect of species richness is more pronounced when greater weight is placed on large biomass values (*q* = 2) than on medium (*q* = 1) or small (*q* = 0.5) values.

### 4.4 Partitioning stability in a metacommunity: assessing spatial synchrony among plots

Because the entire dataset includes 4 blocks and 5 richness levels, we considered all 20 block-richness combinations, resulting in 20 sets of plots. Each set is treated as a metacommunity; see Appendix S4 (Table S4.1) for the distribution of plots across the 20 sets. Each metacommunity comprises 3 or 4 plots/communities, with all plots within a metacommunity containing the same number of species but differing in species composition. For example, the first metacommunity includes 3 plots in Block 1, each with one species; the second one includes 4 plots in Block 1, each with two species; and so on, up to the 20th metacommunity, which includes 3 plots in Block 4, each with 16 species. The biomass in each metacommunity is the sum of biomass values of its constituent plots. We applied stability decomposition to each of the 20 metacommunities. In this context, alpha stability refers to the plot/community level, gamma stability to the metacommunity level (across 3 or 4 plots), and spatial synchrony quantifies the degree of synchrony among plots within each metacommunity.

Based on the complete time series data for plots within each metacommunity, we computed alpha and gamma stability values, as well as among-plot spatial synchrony, specifically for three values of *q* (*q* = 0.5, 1, and 2). For each value of *q*, biodiversity–stability relationships at the gamma and alpha scales, along with biodiversity–synchrony relationships, are presented in Figure 4. For each fixed value of *q*, we fitted a mixed-effects model with random slopes and random intercepts for each block. Figure 4 displays all the fits and significance results.

**Figure 4.**
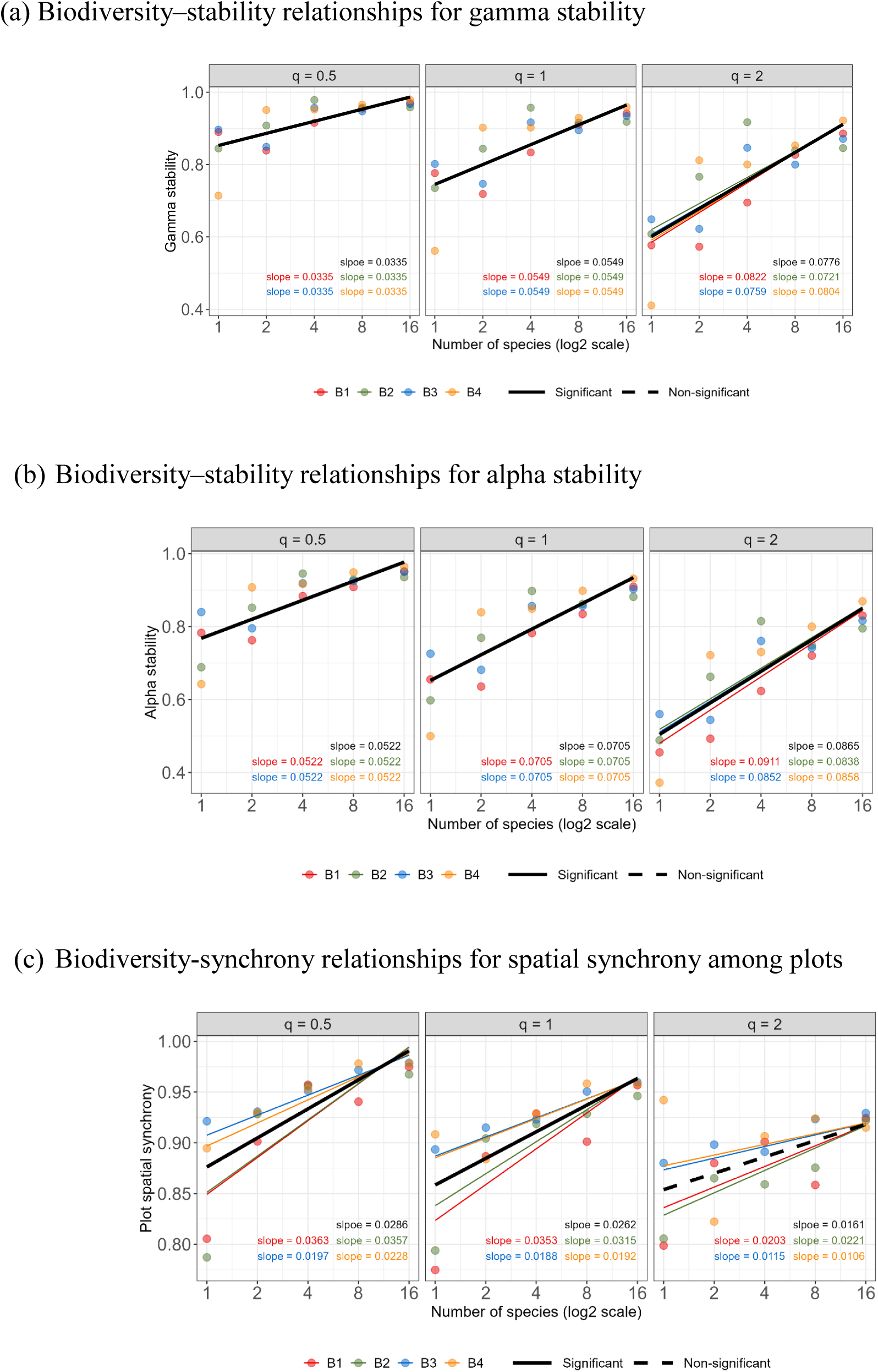
Relationships between the logarithm of species richness and gamma stability (a), alpha stability (b), and spatial synchrony among plots (c) for orders *q* = 0.5, 1 and 2, based on stability decomposition within each of 20 metacommunities. Each metacommunity corresponds to a unique combination of blocks (4 blocks) and species richness level (5 levels); see Table S4.1 (Appendix S4) for plot assignments within each combination. The relationships between species richness and each measure were modeled using a linear mixed-effects model with random slopes and random intercepts for each block. The figure shows overall fixed-effects slopes (bold black lines) and block-specific relationships (thin lines), as estimated from the same linear mixed-effects model. In (a) and (b), because most thin lines nearly overlap with the corresponding bold line, they are not visible. A solid line in the overall fit indicates significance (*P* < 5%), while a dashed line indicates non-significance.

Regardless of alpha stability, gamma stability, or spatial synchrony among plots, positive relationships with biodiversity are consistently found for each fixed value of *q*, with nearly all slopes being significant. These results indicate that increasing species richness promotes community-level stability (Figure 4b), metacommunity-level stability (Figure 4a), and spatial synchrony among plots (Figure 4c). In contrast, Hautier et al. (2020) reached a different conclusion; they found that spatial asynchrony among plots enhanced gamma stability. This discrepancy may be attributed to (i) differences in the stability measures used and (ii) our restriction that local communities have equal species richness. For both gamma and alpha stability, the slope magnitude in Figures 3(a) and 3(b) increases with *q*, suggesting that the effect of species richness is stronger for large biomass than for medium and small biomass.

Figures 2(a) (based on 76 plots) and 4(b) (based on 20 metacommunities) both assess the relationship between biodiversity and plot-level stability, although from different hierarchical decomposition perspectives. Despite the differing approaches, the patterns are consistent, and the fitted slopes are similar in magnitude. In Figure 4(c), for each fixed value of *q*, spatial synchrony is notably low in the two metacommunities composed of monoculture plots. This result indicates that monocultures tend to show greater variation in their relative biomass series compared to mixed species plots.

Unlike the decomposition in Section 4.2, where *asynchrony* among co-occurring species within a plot is the principal determinant of gamma/plot stability, Figure 4 reveals that both alpha stability (Figure 4b) and spatial *synchrony* among plots (Figure 4c) contribute to enhanced meta-community-level gamma stability (Figure 4a). A possible interpretation is that competition among co-occurring species within a plot fosters asynchronous population dynamics, whereas such competition is absent among spatially separated plots (Bessho and Iwasa, 2012; Poethke et al. 2016). This absence allows both alpha stability and spatial *synchrony* to jointly promote gamma stability.

## 5. Discussion

### 5.1 Our choice of information-based distance

As shown in Section 2, we based the temporal information of a biomass time series on the exponential of Rényi entropy, i.e., Hill numbers in the context of species diversity (Eq. 2.1a). However, other classes of information measures also exist. For example, adopting the generalized entropy (also known as Tsallis entropy in physics) leads to alternative formulations. Under this approach, the stability measure for *q* = 1 becomes *H* (***r***) / log *T*, where *H* (***r***) denotes the Shannon entropy based on the relative biomass vector ***r*** = {*r*(*t*); *t* = 1, 2,…,*T*}. For *q* = 2, the corresponding stability measure is 1− *CV* ^2^ (***r***) / *T*. As discussed in Section 2.2, these two measures offer direct and simple ways to take into account the number of time points in time series datasets. Moreover, the decomposition theory presented in Section 3 can also be applied to obtain the associated (a)synchrony measures using these alternative stability formulations.

While these simple measures derived from generalized entropy may seem like promising alternatives, we chose not to adopt them for the following reasons. Suppose the biomass values in a time series are independently identically distributed as a continuous random variable *X* with with probability density function (pdf) *f*(*x*). Let *X* have mean *μ_f_*, variance 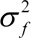, coefficient of variation 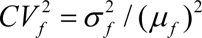 and entropy *H* = *E*(− *X* log *X*) = − (*x* log *x*) *f* (*x*)*dx*. The subscript *f* indicates that these parameters describe properties of the underlying distribution, rather than statistics computed from the observed time series data. As the number of time points increases, it can be shown that both *H* (***r***) / log *T* and 1− *CV* ^2^ (***r***) / *T* tend to 1, regardless of the underlying distribution. As a result, these two simple adjusted measures fail to reflect the true stability or variability of the underlying distribution. In contrast, our measure of *q* = 1 (Eq. 2.3a) converges to a value that reflects stability of the time series via a monotonically increasing function of the Shannon entropy of the random variable *X*. Similarly, our stability measure for *q* = 2 (Eq. 2.3b) converges to 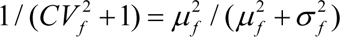, reflecting the stability of the biomass distribution via a monotonically decreasing function of 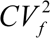. Therefore, our proposed measures accurately characterize the stability of the underlying distribution, whereas the two simple adjusted estimators do not. The derivations of the afore-mentioned limiting values, along with an example in which biomass follows a gamma distribution, appear in Appendix S5.

### 5.2 Comparison of synchrony measures

As indicated in the Introduction, Loreau and de Mazancourt (2008) developed a synchrony measure as a normalized variance of the biomass in the pooled dataset. Since then, the measure has been applied to many studies. Further, Wang and Loreau (2014) obtained the same measure under a variability decomposition framework. In Appendix S3, we provide numerical examples to compare our synchrony measures of *q* = 1 and 2 with this existing measure. The numerical results demonstrate that our synchrony measures offer the following potential improvements over the conventional Loreau and de Mazancourt measure.

1. Sensitivity to data changes: The conventional measure may become insensitive to data variations, whereas our measures respond effectively to changes.
2. Continuity across similar datasets: The conventional measure can yield drastically different values for two slightly different datasets, whereas our measures, in such cases, yield consistently similar synchrony values.
3. Clear quantification target: Our proposed synchrony measure for all *q* > 0 attains a maximum value of 1 if and only if all datasets have identical relative biomass vectors (Section 3.2), providing a clear quantification of similarity among multiple datasets. In contrast, the measure of Loreau and de Mazancourt lacks a well-defined quantification target, as it attains its maximum value of 1 under two distinct conditions: (i) when the correlation between any two datasets is 1, or (ii) when only one dataset fluctuates, while all other datasets show no fluctuation. (In the latter case, correlation between any two datasets is undefined because at least one variance is 0.) Consequently, it does not unambiguously quantify the overall extent of pairwise correlation. See Appendix S3 for examples.

### 5.3 Comparison with a previous analysis of the Jena Experiment data by Roscher et al. (2011)

Using data from the Jena Experiment (2003−2009 and the first harvest only) and the conventional *CV* measure, Roscher et al. (2011) showed that biodiversity enhances community-level stability but reduces species-level population stability. Despite employing different measures from the conventional one, our analysis of the extended Jena dataset (2003−2024) yields results consistent with the findings of Roscher et al. (Figures 2a and 2b). Furthermore, our results show that this opposite effect of biodiversity at the species- and community-level holds not only for large biomass values (*q* = 2) but also for medium and small values within the time series (*q* < 2).

Roscher et al. (2011) also demonstrated that, within a plot, dominant plant species tend to be more stable than intermediate or subordinate/rare species, regardless of species richness. Here, species dominance/rarity is determined by the proportional contribution to total plot biomass, which corresponds to the species relative-biomass weight defined in Section 3.1. In Appendix S6, we plot within-species population stability with respect to species weight. For both orders (*q* = 1 and 2), population stability exhibits a significant positive relationship with species weight, supporting the findings of Roscher et al. They also suggested that lower population stabilities observed at higher species richness can be explained by an increasing number of subordinate species with low stability. In addition, our stability-weight plots in Appendix S6, based on the extended time series data, reveal another insight: some dominant species (e.g., *Onobrychis viciifolia Scop.*, which belongs to the functional group of legumes), are stable at low richness levels but become less stable in highly competitive, high richness environments.

Consistent with many studies that have used the Loreau and de Mazancourt synchrony measure to show that species richness enhances community temporal stability by reducing species synchrony, Figure 2(c) clearly reveals that populations in more-diverse plots tend to be less synchronized than those in less-diverse plots for all values of order *q*. Despite using different measures in our approach, the conclusion remains the same: asynchrony among species plays a key role in promoting the positive biodiversity–stability relationship at the community level.

### 5.4 Extension to hierarchical structure with any number of levels

The stability decomposition theory developed in this paper (Section 3) focuses on a two-level hierarchical structure. For example, in Section 4.1, the stability of each plot/community is decomposed into species-level stability and among-species stability; only two organizational levels (species, community) are involved in the decomposition. Similarly, when assessing spatial synchrony among plots in Section 4.2, the decomposition is also performed at two spatial levels (community, metacommunity). A generalized decomposition framework that integrates both organizational levels and spatial scales into a unified structure was proposed by Wang et al. (2019) using the conventional *CV* measure.

As summarized in Section 4, all plots in the Jena Experiment were deployed in 4 blocks, each containing 18–20 plots positioned at increasing distances from the Saale River near the site (Roscher et al., 2004). While the decomposition framework proposed in this paper focuses on two-level hierarchies, it can be generalized to accommodate more complex structures. For the Jena Experiment data, a multi-level hierarchical decomposition—along with corresponding synchrony measures—can be applied from the entire site level down to individual species:

i. (Block level) The stability of the entire site can be decomposed into within-block (alpha) and among-block (beta) components; the latter beta component is then used to assess spatial (a)synchrony among blocks.
ii. (Plot level) Stability within each block can be decomposed into within- and among-plot stability, allowing for assessment of spatial (a)synchrony among plots.
iii. (Species level) Stability within each plot can be decomposed into within- and among-species population stability, with species-level (a)synchrony also assessable.

Under an additive partitioning scheme, the total stability of the entire site can be expressed as the sum of species-level stability and the three beta components. This approach allows the contribution of each level to total stability to be determined. Details of the extension to accommodate hierarchical structures with more than two levels are currently under development and will be reported in future work.

## 6. Conclusion

Relying on information-based distance/divergence between a temporal data time series and a corresponding maximally stable series (Eq. 2.2a), we have proposed a continuum of variability measures (Eq. 2.2b) and the corresponding stability measures (Eq. 2.2c), and we have applied them to a biomass time series as an example. Both classes of measures are parametrized by an order *q* > 0 and are one-complements of each other. The formulas for the special cases of *q* = 1 (Eq. 2.3a) and *q* = 2 (Eq. 2.3b) are respectively linked to MacArthur’s (1955) entropy-based measure and the conventional *CV* measure. Unlike previous measures, our proposed measures account for the effects of the number of time points over which the measures are computed, enabling meaningful comparisons of ecological data with differing numbers of time points. Numerical examples are given in Section 2.3 and Appendix S1 to demonstrate the importance of incorporating the effect of time. In the special case where biomass remains constant over time (i.e., *CV* = 0), the conventional stability measure (1/*CV* or 1/*CV*^2^) becomes infinite and cannot be further decomposed. In contrast, our stability measure (Eq. 2.3b), by incorporating the number of time points, avoids this infinity issue.

We have also developed a decomposition theory to partition stability across a collection of datasets. The information-based gamma stability (Eq. 3.1a) can be additively or multiplicatively partitioned into an alpha component (Eq. 3.1b) and a beta component. The beta component serves as the basis for deriving mathematically rigorous (a)synchrony measures (Eqs. 3.3b and 3.3c). Importantly, both additive and multiplicative schemes lead to the same measures of (a)synchrony. In Appendix S3, we compare our synchrony measures with the measure developed by Loreau and de Mazancourt (2008). Our theoretical analysis and numerical results suggest that our synchrony measures establish a mathematically rigorous formulation and offer potential improvements over previous approaches.

In addition to examining the effects of biodiversity on temporal stability, the relationships between biodiversity and spatial stability have also been widely explored (Weigelt et al., 2008; Eisenhauer et al., 2011; Gottschall et al., 2022). For example, Weigelt et al. (2008) found that functional diversity, as measured by Rao’s quadratic entropy, decreases spatial variability (as quantified by the *CV*) among subplots within plots. We affirm that our proposed continuum of stability measures and decomposition framework can also be applied to assess spatial stability and variability for any non-negative variables.

## Acknowledgements

This work is supported by the Taiwan National Science and Technology Council under Contracts NSTC-113-2118-M-007-005 (for AC) and by BETA-FOR (for ST, OM, BMD, AF, NE, and JM). BETA-FOR is funded by the 12249 (DFG, German Research Foundation, FOR 5375) - 459717468, with additional support by the Julius-Maximilians-Universität Würzburg (JMU). NE acknowledges support from the German Centre for Integrative Biodiversity Research Halle– Jena–Leipzig, funded by the German Research Foundation (DFG; FZT 118, 202548816), as well as by the DFG (Ei 862/29-1).

## CONFLICT OF INTEREST STATEMENT

The authors declare no conflict of interest.

## Notes

https://github.com/AnneChao/MS_iSTAY

### Competing Interest Statement

The authors have declared no competing interest.

