## Appendices for "A continuum of information-based temporal stability measures and their decomposition across hierarchical levels"

#### Appendix S1. Mathematical proofs and examples

*The coefficient of variation (CV) is primarily sensitive to large data values in a biomass time series*

Let $\{z(t); t = 1, 2, ..., T\}$ denote the original biomass time series. The corresponding relative biomass vector is denoted by $\boldsymbol{r} = \{r(t); t = 1, 2, ..., T\}$ with a mean of $1/T$, where $r(t) = z(t)/z$ and $z = z(+) = \sum_{t=1}^{T} z(t)$. The $CV^2$ of the original biomass vector is equal to that of the relative biomass vector and thus can be expressed as:

$$CV^2(\boldsymbol{r}) = \frac{\mathrm{var}(\boldsymbol{r})}{(mean(\boldsymbol{r}))^2} = \frac{\frac{1}{T-1}\sum_{t=1}^{T}[r(t) - 1/T]^2}{(1/T)^2} = \frac{T^2}{T-1}\sum_{t=1}^{T}[r(t)]^2 - \frac{T}{T-1}. \qquad \text{(S1.1)}$$

In the expression above, $\sum_{t=1}^{T}[r(t)]^2$ represents the Simpson index based on the relative abundance vector $\boldsymbol{r} = \{r(t); t = 1, 2, ..., T\}$. As a result, larger biomass values are given disproportionally more weights; the $CV$ is therefore primarily sensitive to large data values.

*The maximum value that $CV^2$ can attain, for T time points, is T*

__Theorem S1.1__ **(The range of $CV^2$ of a biomass series for $T$ time points is [0, $T$])** The maximum value that the $CV^2$ of a biomass time series can attain over $T$ time points is $T$. Here the maximum is obtained by considering all possible relative vectors with $T$ components.

**Proof**: For biomass data collected from $T$ time points, the term $\sum_{t=1}^{T}[r(t)]^2$ in Eq. (S1.1) attains the maximum value of 1 if one value dominates, i.e., one relative biomass value is close to 1, and the other $T-1$ values are close to 0. In this extreme case, $\sum_{t=1}^{T}[r(t)]^2 \to 1$. Then the maximum value that $CV^2$ can attain is $T$. QED

*The importance of incorporating time series length in stability measures*

In the main text (Section 2.3), a numerical example was provided to intuitively explain why the effect of time series length should be considered in the conventional $CV^2$ measure (*i.e.*, $q$ = 2). We explained that two hypothetical biomass time series (referred to as Series A and B in Section 2.3) have the same value of $CV$, but intuitively, the two series should not be equally stable. Based on our stability measures, the stability profiles for the two series are shown in Figure S1.1 (a). For any $q$ between 0 and 2, Series A is consistently less stable than Series B, to an extent dependent on $q$. In the special case of $q = 2$, our proposed measure for Series A is 0.002, whereas for Series B it is 0.286.

We now give a numerical example for $q = 1$ to further illustrate the importance of incorporating time series length. In Section 2.2 of the main text, we explained why Shannon entropy, in its traditional form, cannot be applied—alone—to quantify stability, and why time series length should also be considered. That is, instead of entropy $H(\boldsymbol{r})$ (Eq. 2.1c in the main text), our proposed measure (Eq. 2.3a in the main text) is $\{\exp[H(\boldsymbol{r})]-1\}/(T-1)$, where $T$ denotes time series length. Consider a biomass Series C (time series length = 3), in which the three relative biomass values are {0.25, 0.3, 0.45}. For this series, we have Shannon entropy $H(\boldsymbol{r}) = 1.067$. Consider another time series, Series D (time series length = 19), with relative biomass values given by {1 × 0.8, 2 × 0.2, 16 × 0.01} —comprising one dominant value (0.8), two small values (0.2), and sixteen very small values (0.01). Intuitively, Series C appears much more stable than Series D. However, Series D yields an entropy value of 1.072, nearly identical to that of Series C, leading to the misleading implication that the two series are equally stable.

The issue described above arises from the use of Shannon entropy because, for $T = 3$, the maximum possible value of $\exp[H(\boldsymbol{r})]-1$ is $T - 1 = 2$, which occurs when biomass remains constant over all three time points. Accordingly, Series C (with $\exp[H(\boldsymbol{r})]-1 =$ 1.907) represents a highly stable series. When $T = 19$, the maximum value is increased to 18 (when biomass remains constant over all 19 time points). In this case, Series D (with $\exp[H(\boldsymbol{r})]-1$ = 1.921) reflects a much lower degree of stability. Based on our proposed measures, the stability for Series C is $\{\exp[H(\boldsymbol{r})]-1\}/(T-1)$ = 1.907/2 = 0.9535 (variability = 0.0465), whereas the stability for Series D is 1.921/18 = 0.107 (variability = 0.893), clearly signifying that Series C is more stable than Series D. As shown in Figure S1.1(b), this conclusion holds for any $q > 0$ under our stability measures.

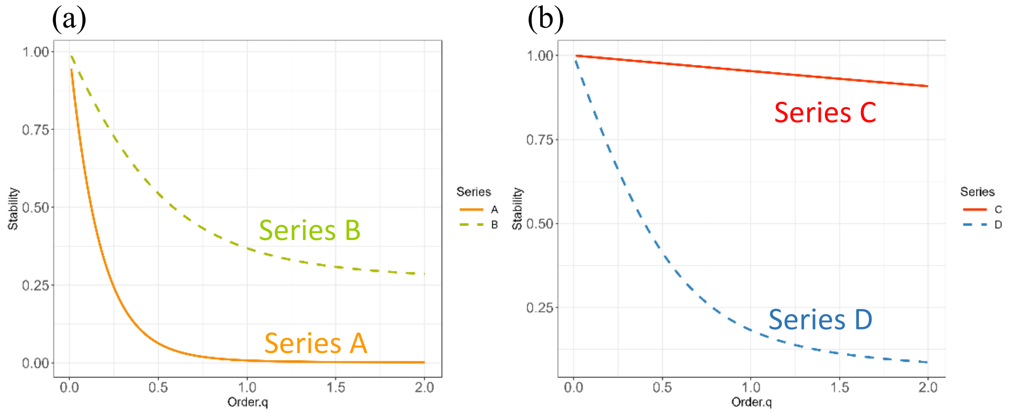


**<u>Figure S1.1</u> Stability profiles of our measures for order $q$ between 0 and 2.** (a) The relative biomass sets for Series A and B are {0.999, 0.001} ($T = 2$) and {3 × 0.3049, 6 × 0.0122, 1 × 0.0121} ($T = 10$), respectively; see main text for details. The two series yield nearly identical $CV$ values, but the profiles of our proposed stability measures indicate that Series B is more stable than Series A. (b) The relative biomass distributions for Series C and D are {0.25, 0.3, 0.45} ($T = 3$) and {1 × 0.8, 2 × 0.2, 16 × 0.01} ($T = 19$), respectively. The two series have nearly equal entropy values, but the profiles of our stability measures clearly demonstrate that Series C is substantially more stable than Series D.

##### *Inequalities between alpha and gamma stability, and the maximum values of beta stability measures*

As derived in Section 3 of the main text, we have the following formulas for gamma and alpha stability:

Gamma stability:
$$ {}^qS_\gamma = \begin{cases} \dfrac{{}^qI(\mathbf{r}^{(\gamma)})-1}{T-1} = \dfrac{1}{T-1}\left\{\left[\sum_{t=1}^{T}\left(\dfrac{z_+(t)}{z_+}\right)^q\right]^{\frac{1}{1-q}}-1\right\}, & q\neq 1\\[2ex] \dfrac{1}{T-1}\left\{\exp\left[-\sum_{t=1}^{T}\left(\dfrac{z_+(t)}{z_+}\right)\log\left(\dfrac{z_+(t)}{z_+}\right)\right]-1\right\}, & q=1 \end{cases} $$

Alpha stability:
$$ {}^qS_\alpha = \begin{cases} \dfrac{\left[\sum_{k=1}^{K} w_k[{}^qI(\mathbf{r}^{(k)})]^{1-q}\right]^{1/(1-q)}-1}{T-1} = \dfrac{1}{T-1}\left[\left\{\sum_{k=1}^{K} w_k\left[\sum_{t=1}^{T}\left(\dfrac{z_k(t)}{z_k}\right)^q\right]\right\}^{\frac{1}{1-q}}-1\right], & q\neq 1\\[2ex] \dfrac{1}{T-1}\left[\exp\left\{-\sum_{k=1}^{K} w_k\left[\sum_{t=1}^{T}\left(\dfrac{z_k(t)}{z_k}\right)\log\left(\dfrac{z_k(t)}{z_k}\right)\right]\right\}-1\right], & q=1 \end{cases} $$

In the following two theorems, we prove that gamma stability is never less than alpha stability for the general case $q > 0$, $q \neq 1$ (Theorem S1.2) and for the special case of $q = 1$ (Theorem S1.3). In each case, we also derive a data-dependent maximum of beta stability when alpha stability is fixed. That is, the maximum (denoted as $\max({}^qS_\beta)$ in Theorem S1.2 and $\max({}^1S_\beta)$ in Theorem S1.3) is obtained given the collection of $K$ biomass vectors of individual datasets; see Appendix S2 for further numerical illustration.

**Theorem S1.2 (for general $q > 0$, $q \neq 1$).** For the proposed gamma and alpha stability measures, we have the following inequalities for $q > 0$ and $q \neq 1$:

$$ 0 \le {}^qS_\gamma - {}^qS_\alpha \le \frac{\max\ {}^qI(\mathbf{r}^{(\gamma)})-1}{T-1} - {}^qS_\alpha \equiv \max({}^qS_\beta). $$

Here $\max {}^qI(\boldsymbol{r}^{(\gamma)}) = \min\{{}^qI(\boldsymbol{w}\times\boldsymbol{r}), (1-1/K)T + (1/K)[{}^qI(\boldsymbol{w}\times\boldsymbol{r})]\}$ and ${}^qI(\boldsymbol{w}\times\boldsymbol{r})$ denotes the information computed from the two-dimensional $K$ by $T$ table $[w_k r_k(t)]$, i.e.,

$$ {}^qI(\boldsymbol{w}\times\boldsymbol{r}) = \begin{cases} \left[\sum_k\sum_t [w_k r_k(t)]^q\right]^{1/(1-q)}, & q \neq 1 \\ \exp[H(\boldsymbol{w}\times\boldsymbol{r})], & q = 1 \end{cases} $$

where $H(\boldsymbol{w}\times\boldsymbol{r}) = -\sum_k\sum_t [w_k r_k(t)]\log[w_k r_k(t)]$ denotes Shannon entropy based on $[w_k r_k(t)]$.

The minimum value of 0 for beta stability is attained (i.e., ${}^qS_\gamma = {}^qS_\alpha$) if and only if all datasets have identical relative biomass time series. The above maximum is attained if and only if there exists a dataset $\ell$ such that $z_+(t) = z_\ell(t)$. That is, at any time point, the total biomass in the pooled dataset is almost exclusively contributed from only one dataset, while the other datasets have negligible biomass.

**Proof**:

For $q > 1$, $f(x) = x^q$ is a convex function, and $\frac{z_+(t)}{z_+} = \sum_{k=1}^{K} \frac{z_k}{z_+}\frac{z_k(t)}{z_k}$, it follows from the Jensen inequality that for any fixed time $t$, we have

$$ \left(\frac{z_+(t)}{z_+}\right)^q = \left(\sum_{k=1}^{K} \frac{z_k}{z_+}\frac{z_k(t)}{z_k}\right)^q \leq \sum_{k=1}^{K} \frac{z_k}{z_+}\left(\frac{z_k(t)}{z_k}\right)^q. $$

Summing over all time points, we obtain ${}^qS_\gamma - {}^qS_\alpha \geq 0$. The Jensen inequality becomes an equality if and only if $\frac{z_1(t)}{z_1} = \frac{z_2(t)}{z_2} = \ldots = \frac{z_K(t)}{z_K}$ for any fixed time $t$ = 1, 2, …, $T$, i.e., all $K$ datasets have identical biomass relative distributions. For $q < 1$, $f(x) = x^q$ is, instead, a concave function, a similar proof also leads to ${}^qS_\gamma - {}^qS_\alpha \geq 0$. The maximum value of ${}^qS_\gamma - {}^qS_\alpha$ is obtained through the following derivation:

$$ {}^qS_\gamma - {}^qS_\alpha = \frac{{}^qI(\boldsymbol{r}^{(\gamma)}) - 1}{T-1} = \frac{1}{T-1}\left\{\left[\sum_{t=1}^{T}\left(\frac{z_+(t)}{z_+}\right)^q\right]^{\frac{1}{1-q}} - 1\right\} - {}^qS_\alpha $$

$$ \leq \frac{1}{T-1}\left\{\left[\sum_{t=1}^{T}\sum_{k=1}^{K}\left(\frac{z_k(t)}{z_+}\right)^q\right]^{\frac{1}{1-q}} - 1\right\} - {}^qS_\alpha . $$

$$ = \frac{{}^qI(\boldsymbol{w}\times\boldsymbol{r}) - 1}{T-1} - {}^qS_\alpha $$

However, ${}^qI(\boldsymbol{w}\times\boldsymbol{r})$ is generally increases rapidly with $K$ and thus cannot be used as a tight upper bound. Note that ${}^qI(\boldsymbol{r}^{(\gamma)}) \leq T$, implying that any convex combination of ${}^qI(\boldsymbol{w}\times\boldsymbol{r})$ and

$T$ can be used as an appropriate upper bound of ${}^qI(\boldsymbol{r}^{(\gamma)})$. We choose a tight upper bound as follows:

$$\max {}^qI(\boldsymbol{r}^{(\gamma)}) = \begin{cases} {}^qI(\boldsymbol{w}\times\boldsymbol{r}) & if\ {}^qI(\boldsymbol{w}\times\boldsymbol{r}) \le T \\ (1-\dfrac{1}{K})T + \dfrac{1}{K}[{}^qI(\boldsymbol{w}\times\boldsymbol{r})] & if\ {}^qI(\boldsymbol{w}\times\boldsymbol{r}) > T \end{cases}$$

Equivalently, $\max {}^qI(\boldsymbol{r}^{(\gamma)}) = \min\{{}^qI(\boldsymbol{w}\times\boldsymbol{r}), (1-1/K)T + (1/K)[{}^qI(\boldsymbol{w}\times\boldsymbol{r})]\}$. Our final inequality becomes an equality if and only if ${}^qI(\boldsymbol{r}^{(\gamma)}) = {}^qI(\boldsymbol{w}\times\boldsymbol{r})$, i.e., there exists a dataset $\ell$ such that $z_+(t) = z_\ell(t)$. That is, at any time point, the total biomass in the pooled dataset is almost exclusively contributed from only one dataset, while the other datasets have negligible biomass. QED

**<u>Theorem S1.3 (for *q* = 1).</u>** Given that alpha stability is fixed, we have the following inequality for the corresponding entropy-based gamma and alpha stability measures:

$$0 \le {}^1S_\gamma - {}^1S_\alpha \le \frac{\max {}^1I(\boldsymbol{r}^{(\gamma)}) - 1}{T-1} - {}^qS_\alpha \equiv \max({}^1S_\beta).$$

Here $\max {}^1I(\boldsymbol{r}^{(\gamma)}) = \min\{{}^1I(\boldsymbol{w}\times\boldsymbol{r}), (1-1/K)T + (1/K)[{}^1I(\boldsymbol{w}\times\boldsymbol{r})]\}$, ${}^1I(\boldsymbol{w}\times\boldsymbol{r}) = \exp[H(\boldsymbol{w}\times\boldsymbol{r})]$, and $H(\boldsymbol{w}\times\boldsymbol{r}) = -\sum_k\sum_t [w_k r_k(t)]\log[w_k r_k(t)]$ denotes Shannon entropy based on the values in the matrix $[w_k r_k(t)]$. The minimum value of 0 is attained (i.e., ${}^1S_\gamma = {}^1S_\alpha$) if and only if all datasets have identical relative biomass time series. The maximum is attained if and only if, at any time point, the total biomass in the pooled dataset is almost exclusively contributed from only one dataset, while the other datasets have negligible biomass.

**Proof**:

Since $f(x) = -x\log x$ is a concave function and $\dfrac{z_+(t)}{z_+} = \sum_{k=1}^{K}\dfrac{z_k}{z_+}\dfrac{z_k(t)}{z_k}$, it follows from the Jensen inequality that for any fixed time *t*, we have

$$-\frac{z_+(t)}{z_+}\log\left(\frac{z_+(t)}{z_+}\right) \ge -\sum_{k=1}^{K}\frac{z_k}{z_+}\frac{z_k(t)}{z_k}\log\frac{z_k(t)}{z_k}.$$

Summing over all time points, we obtain

$$-\sum_{t=1}^{T}\frac{z_+(t)}{z_+}\log\left(\frac{z_+(t)}{z_+}\right) \ge -\sum_{k=1}^{K}\frac{z_k}{z_+}\sum_{t=1}^{T}\frac{z_k(t)}{z_k}\log\frac{z_k(t)}{z_k}$$

This proves ${}^1S_\gamma \ge {}^1S_\alpha$. The Jensen inequality become an equality if and only if $\dfrac{z_1(t)}{z_1} = \dfrac{z_2(t)}{z_2} = \ldots = \dfrac{z_K(t)}{z_K}$ for any fixed time *t* = 1, 2, …, *T*, i.e., all *K* datasets have identical biomass relative distributions. The maximum value of ${}^1S_\gamma - {}^1S_\alpha$ is obtained through the following derivation:

$$
\begin{aligned}
{}^1S_\gamma - {}^1S_\alpha &= \frac{\exp\left[H(r^{(\gamma)})\right]-1}{T-1} - {}^1S_\alpha \\
&= \frac{\exp\left[-\sum_{t=1}^{T}\frac{z_+(t)}{z_+}\log\frac{z_+(t)}{z_+}\right]-1}{T-1} - {}^1S_\alpha \\
&\le \frac{\exp\left[-\sum_{t=1}^{T}\left(\sum_{k=1}^{K}\frac{z_k}{z_+}\frac{z_k(t)}{z_k}\right)\log\left(\frac{z_k}{z_+}\frac{z_k(t)}{z_k}\right)\right]-1}{T-1} - {}^1S_\alpha \\
&= \frac{{}^1I(\boldsymbol{w}\times\boldsymbol{r})-1}{T-1} - {}^1S_\alpha
\end{aligned}
$$

The proof is then completed based on similar procedures as those for a general order $q$. QED

Based on Theorems S1.2 and S1.3, our measures of asynchrony (Eqs. 3.3a and 3.3b in the main text) and synchrony (Eq. 3.3c) among datasets turn out to be

$$
\begin{aligned}
{}^q\overline{\psi} &= \frac{{}^qS_\beta}{\max({}^qS_\beta)} \\
&= \frac{{}^qI(\boldsymbol{r}^{(\gamma)}) - \left[\sum_{k=1}^{K} w_k[{}^qI(\boldsymbol{r}^{(k)})]^{1-q}\right]^{1/(1-q)}}{\max\ {}^qI(\boldsymbol{r}^{(\gamma)}) - \left[\sum_{k=1}^{K} w_k[{}^qI(\boldsymbol{r}^{(k)})]^{1-q}\right]^{1/(1-q)}}, \quad q \ne 1.
\end{aligned}
$$

$$
{}^q\psi = 1 - {}^q\overline{\psi} = \frac{\max\ {}^qI(\boldsymbol{r}^{(\gamma)}) - {}^qI(\boldsymbol{r}^{(\gamma)})}{\max\ {}^qI(\boldsymbol{r}^{(\gamma)}) - \left[\sum_{k=1}^{K} w_k[{}^qI(\boldsymbol{r}^{(k)})]^{1-q}\right]^{1/(1-q)}}, \quad q \ne 1.
$$

#### Appendix S2. Use numerical examples to illustrate how the maximum values in Eq. (3.2a) is obtained

Given the collection of $K$ biomass vectors of individual datasets, a data-dependent maximum for beta stability of any order $q > 0$ is derived in Appendix S1. The resulting maximum is also stated in Eq. (3.2a) of the main text. In this Appendix, two simple examples (Examples 1 and 2) are used to illustrate how the maximum is obtained. In Example 1, the maximum can be attained by a dataset, whereas in Example 2 the maximum cannot be attained by any dataset.

**Example 1** **(The maximum of $^{q}S_{\beta} = {}^{q}S_{\gamma} - {}^{q}S_{\alpha}$ can be attained)** Consider an original collection of two datasets with time series length = 4 (columns): the first dataset consists of a biomass time series [3 2 0 0] and the second data set consists of another time series [0 6 0 4]. We represent this original collection as the matrix $\begin{bmatrix} 3 & 2 & 0 & 0 \\ 0 & 6 & 0 & 4 \end{bmatrix}$ in Table S2.1, i.e., the two datasets are displayed as two rows. Within each dataset, we can permute biomass values without affecting its stability, as stability is independent of time ordering. For example, the matrix $\begin{bmatrix} 3 & 0 & 2 & 0 \\ 0 & 6 & 0 & 4 \end{bmatrix}$ has the same alpha stability as the original matrix. However, the two matrices have different gamma values and thus exhibit different beta stability, as the pooled datasets differ from each other.

Given that alpha stability is fixed—given the two data series—there are several matrices that give the same value of alpha stability. We list five other matrices in the following table, all of which have the same alpha stability as the original data matrix but exhibit different gamma stability, leading to varying beta stability. For each matrix, the beta stability and its maximum value are shown. Each maximum (0.581 for $q = 1$ and 0.513 for $q = 2$) represents the theoretical upper bound among all possible matrices with a fixed alpha stability. For this example, the maximum is attained by the matrix (shown in red) where the total biomass in the pooled dataset at any time point is primarily contributed by only one dataset, while the other dataset has negligible biomass. This means that in each column, there is only one positive value, and the other values are close to 0.

**<u>Table S2.1</u>**. Alpha and beta stability for $q$ = 1 and $q$ = 2, and the attainable maximum value of beta stability for six matrices that have the same alpha value. The original matrix is shown in blue; the matrix shown in red attains the maximum beta stability.

<table>
<tr><th rowspan="2">Matrix</th><th colspan="3">$q = 1$</th><th colspan="3">$q = 2$</th></tr>
<tr><th>${}^{1}S_{\alpha}$</th><th>${}^{1}S_{\beta} = {}^{1}S_{\gamma} - {}^{1}S_{\alpha}$</th><th>$\max({}^{1}S_{\beta})$</th><th>${}^{2}S_{\alpha}$</th><th>${}^{2}S_{\beta} = {}^{2}S_{\gamma} - {}^{2}S_{\alpha}$</th><th>$\max({}^{2}S_{\beta})$</th></tr>
<tr><td>$\begin{bmatrix} 3 & 0 & 2 & 0 \\ 0 & 6 & 0 & 4 \end{bmatrix}$</td><td rowspan="6">0.320</td><td>0.581</td><td rowspan="6">0.581</td><td rowspan="6">0.308</td><td>0.513</td><td rowspan="6">0.513</td></tr>
<tr><td>$\begin{bmatrix} 3 & 0 & 2 & 0 \\ 0 & 6 & 4 & 0 \end{bmatrix}$</td><td>0.304</td><td>0.285</td></tr>
<tr><td>$\begin{bmatrix} 3 & 2 & 0 & 0 \\ 0 & 6 & 0 & 4 \end{bmatrix}$</td><td>0.261</td><td>0.202</td></tr>
<tr><td>$\begin{bmatrix} 2 & 0 & 3 & 0 \\ 0 & 6 & 4 & 0 \end{bmatrix}$</td><td>0.244</td><td>0.202</td></tr>
<tr><td>$\begin{bmatrix} 3 & 2 & 0 & 0 \\ 4 & 6 & 0 & 0 \end{bmatrix}$</td><td>0.012</td><td>0.023</td></tr>
<tr><td>$\begin{bmatrix} 2 & 3 & 0 & 0 \\ 4 & 6 & 0 & 0 \end{bmatrix}$</td><td>0</td><td>0</td></tr>
</table>

**<u>Example 2</u> (The maximum of ${}^{q}S_{\beta} = {}^{q}S_{\gamma} - {}^{q}S_{\alpha}$ cannot be attained)** Consider an original collection of two datasets (rows) with time series length = 3 (columns): the first dataset consists of a biomass time series [3 0 2], and the second dataset consists of another biomass time series [1 5 0]. We display this collection of two datasets as the matrix $\begin{bmatrix} 3 & 0 & 2 \\ 1 & 5 & 0 \end{bmatrix}$ in Table S2.2.

As with Example 1, given that alpha stability is fixed, we list five other matrices in the following table; all have the same alpha stability as the original matrix but exhibit different gamma and beta stability. For example, regardless of order $q$, the first matrix has higher gamma stability and, consequently, higher beta stability than the original matrix. For each matrix, the beta stability and its maximum value for $q$ = 1 and 2 are shown. Each maximum (0.861 for $q$ = 1 and 0.758 for $q$ = 2) represents the theoretical upper bound among all possible matrices with a fixed alpha stability. For this example, the maximum cannot be attained for any $q$ > 0 because the condition that “the total biomass in the pooled dataset, at any time point, is primarily contributed by only one dataset” cannot be met under the condition of fixed alpha stability. Nevertheless, the theoretical maximum is still valid for normalization purposes, although it is not attainable.

**Table S2.2**. Alpha and beta stability of $q = 1$ and $q = 2$, and the unattainable maximum value of beta stability for six matrices that have the same alpha value. The original matrix is shown in blue.

| Matrix | $q=1$ | | | $q=2$ | | |
|---|---|---|---|---|---|---|
| | ${}^{1}S_{\alpha}$ | ${}^{1}S_{\beta} = {}^{1}S_{\gamma} - {}^{1}S_{\alpha}$ | $\max({}^{1}S_{\beta})$ | ${}^{2}S_{\alpha}$ | ${}^{2}S_{\beta} = {}^{2}S_{\gamma} - {}^{2}S_{\alpha}$ | $\max({}^{2}S_{\beta})$ |
| $\begin{bmatrix}3&0&2\\0&5&1\end{bmatrix}$ | 0.368 | 0.585 | 0.746 | 0.293 | 0.614 | 0.732 |
| $\begin{bmatrix}3&0&2\\1&5&0\end{bmatrix}$ | | 0.541 | | | 0.551 | |
| $\begin{bmatrix}3&0&2\\0&1&5\end{bmatrix}$ | | 0.313 | | | 0.232 | |
| $\begin{bmatrix}3&0&2\\5&1&0\end{bmatrix}$ | | 0.201 | | | 0.084 | |
| $\begin{bmatrix}3&0&2\\1&0&5\end{bmatrix}$ | | 0.095 | | | 0.138 | |
| $\begin{bmatrix}3&0&2\\5&0&1\end{bmatrix}$ | | 0.030 | | | 0.035 | |

#### Appendix S3. Comparisons of synchrony measures

Following the framework in Section 3 of the main text, we consider a collection of $K$ datasets, each representing a time series of biomass or another pertinent variable. The data from each set can be obtained from sampling units at different levels of ecological organization (e.g., species, guilds, functional groups, habitats, communities, or ecosystems) or at various spatial scales (e.g., local, regional, or any spatial unit). Our proposed synchrony of a general order $q > 0$ (Eq. 3.3c) measures the extent of similarity or closeness among the $K$ relative datasets. In this Appendix, we focus on comparing our measures of $q = 1$ and 2 with the conventional synchrony measure developed by Loreau and de Mazancourt (2008).

For simplicity, the synchrony measure developed by Loreau and de Mazancourt (2008) is denoted as $\phi_{\mathrm{LM}}$, and our proposed measures are denoted as ${}^1\psi$ (for $q = 1$) and ${}^2\psi$ (for $q = 2$). The formula for ${}^q\psi$ for a general order $q > 0$ is shown in Eq. (3.3c); the formula specifically for ${}^1\psi$ is given in Eq. (3.5b) of the main text. Based on the notation in Section 3 of the main text, we let $z_k(t)$ denote the biomass, productivity or another variable for dataset $k$ at time point $t$, $k = 1, 2, \ldots, K$ and $t = 1, 2, \ldots, T$. Dataset $k$ consists of the time series $\{z_k(t); t = 1, 2, \ldots, T\}$. As in Appendix S2, we represent the collection of $K$ datasets as a $K \times T$ dimensional matrix $[z_k(t)]$. For example, if we consider a collection of two datasets (rows) with time series length = 4 (columns), we represent this collection of two datasets as the matrix $\begin{bmatrix} z_1(1) & z_1(2) & z_1(3) & z_1(4) \\ z_2(1) & z_2(2) & z_2(3) & z_2(4) \end{bmatrix}$. To intuitively compare $\phi_{\mathrm{LM}}$ with our measures, we mainly focus on collections of two or three datasets (rows, $K = 2$ or 3) with a time series length of three or four (columns, $T = 3$ or 4). All comparisons can be extended to any arbitrary numbers of datasets and time series lengths.

Loreau and de Mazancourt (2008) originally developed the measure $\phi_{\mathrm{LM}}$ as a normalized variance of the biomass in the pooled dataset. Let $\sigma_{jk}$ denote the covariance between dataset $k$ and dataset $j$. If $k = j$, then $\sigma_{kk}$ becomes the variance of dataset $k$. The formula for $\phi_{\mathrm{LM}}$ can be expressed as

$$\phi_{LM} = \frac{Var\left(\sum_{k=1}^{K} z_k(t)\right)}{\max\left\{Var\left(\sum_{k=1}^{K} z_k(t)\right)\right\}} = \frac{\sum_{k=1}^{K}\sum_{j=1}^{K}\sigma_{jk}}{\left[\sum_{k=1}^{K}\sqrt{\sigma_{kk}}\right]^2}. \qquad \text{(S3.1)}$$

Wang and Loreau (2014) derived the same synchrony measure based on the variability decomposition framework developed in Thibaut and Connolly (2013). The numerator in Eq. (S3.1) represents the variance of the summed biomass over datasets, and the denominator represents the maximum value of the variance.

For the special case of $K = 2$, the measure $\phi_{\mathrm{LM}}$ reduces to

$$\phi_{LM} = \frac{\sigma_{11} + \sigma_{22} + 2\sigma_{12}}{\sigma_{11} + \sigma_{22} + 2\sqrt{\sigma_{11}\sigma_{22}}} = \frac{\text{var}_1 + \text{var}_2 + 2\,\text{cov}_{12}}{\text{var}_1 + \text{var}_2 + 2\sqrt{\text{var}_1\ \text{var}_2}}. \tag{S3.2}$$

Here $\text{var}_1$, $\text{var}_2$ and $\text{cov}_{12}$ denote the variance of dataset 1, variance of dataset 2, and the covariance between dataset 1 and dataset 2 respectively. The measure attains the maximum value of 1 if $\text{cov}_{12} = \sqrt{\text{var}_1\ \text{var}_2}$ , i.e., the correlation between the two datasets is 1, if both variances > 0. However, if one of the two variances is zero, say $\text{var}_1 = 0$, then $\text{cov}_{12} = 0$; $\text{cov}_{12} = \sqrt{\text{var}_1\ \text{var}_2}$ is also satisfied and $\phi_{\text{LM}} = 1$ for *any* dataset 2. This leads to two interpretational problems, as discussed below. Similar problems arise when there are more than two datasets. Note that a *prerequisite for the measure* $\phi_{\text{LM}}$ to make sense is the existence of at least one dataset with variance greater than zero. Otherwise, the denominators in both Eqs. (S3.1) and (S3.2) become zero, rendering the measure $\phi_{\text{LM}}$ undefined.

(1) *The measure $\phi_{LM}$ may become insensitive to data*

In the special case of $K = 2$, as stated earlier, if there is no fluctuation in the first dataset ($\text{var}_1 = 0$ in Eq. S3.2), both the denominator and numerator are equal to $\text{var}_2$. Then, the measure $\phi_{LM}$ always attains the maximum value of 1, regardless of the values in the second dataset. The measure $\phi_{\text{LM}}$ becomes insensitive to the data of the second dataset. That is, consider the matrix $\begin{bmatrix} c & c & c & c \\ x_1 & x_2 & x_3 & x_4 \end{bmatrix}$ where $c$ denotes a constant. Regardless of the values of $x_i$, this matrix always attains the maximum synchrony value of 1. By contrast, our proposed measures are sensitive to changes in the second dataset, as revealed by the three examples below.

| Biomass matrix (row: dataset column: time) | | $A_1 = \begin{bmatrix} 5 & 5 & 5 & 5 \\ 5 & 5 & 5 & 4 \end{bmatrix}$ | $A_2 = \begin{bmatrix} 5 & 5 & 5 & 5 \\ 8 & 6 & 9 & 3 \end{bmatrix}$ | $A_3 = \begin{bmatrix} 5 & 5 & 5 & 5 \\ 8 & 60 & 9 & 30 \end{bmatrix}$ |
|---|---|---|---|---|
| Loreau and de Mazancourt (2008) | | 1 | 1 | 1 |
| Proposed | $q = 1$ | 0.998 | 0.964 | 0.893 |
| | $q = 2$ | 0.996 | 0.943 | 0.806 |

Similar insensitivity arises when there are more than two datasets in a matrix. Consider the following three matrices, in each of which the first two datasets remain constant over time. Consequently, the measure $\phi_{\text{LM}}$ always attains the maximum value of 1, regardless of the biomass values in the third dataset. That is, if we have the matrix $\begin{bmatrix} c & c & c & c \\ c' & c' & c' & c' \\ x_1 & x_2 & x_3 & x_4 \end{bmatrix}$ where $c$ and $c'$ denote two constants, then regardless of the values of $x_i$, this matrix always attains the maximum synchrony value of 1. By contrast, our proposed measures are sensitive to changes in the third dataset. See the three examples below.

| Biomass matrix (row: dataset column: time) | | $B_1 = \begin{bmatrix} 2 & 2 & 2 & 2 \\ 3 & 3 & 3 & 3 \\ 1 & 4 & 5 & 8 \end{bmatrix}$ | $B_2 = \begin{bmatrix} 2 & 2 & 2 & 2 \\ 3 & 3 & 3 & 3 \\ 11 & 14 & 15 & 18 \end{bmatrix}$ | $B_3 = \begin{bmatrix} 2 & 2 & 2 & 2 \\ 3 & 3 & 3 & 3 \\ 10 & 40 & 50 & 80 \end{bmatrix}$ |
|---|---|---|---|---|
| Loreau and de Mazancourt (2008) | | 1 | 1 | 1 |
| Proposed | $q = 1$ | 0.927 | 0.992 | 0.926 |
| | $q = 2$ | 0.887 | 0.977 | 0.889 |

*(2) The measure $\phi_{LM}$ may give drastically different values for slightly different matrices*

In the following table, we give three matrices that differ slightly only at the last time point. However, the synchrony value $\phi_{LM}$ for the first matrix $C_1$ is minimally zero (because the biomass in the pooled set is a constant), the synchrony value for $C_2$ is maximally unity (because there is only one dataset with fluctuation), and the synchrony value for $C_3$ takes a value in between. For each $q = 1$ or 2, the synchrony values of our measures are slightly different, but all are close to 1.

| Biomass matrix (row: dataset column: time) | | $C_1 = \begin{bmatrix} 5 & 5 & 5 & 6 \\ 5 & 5 & 5 & 4 \end{bmatrix}$ | $C_2 = A_1 = \begin{bmatrix} 5 & 5 & 5 & 5 \\ 5 & 5 & 5 & 4 \end{bmatrix}$ | $C_3 = \begin{bmatrix} 5 & 5 & 5 & 5.5 \\ 5 & 5 & 5 & 3 \end{bmatrix}$ |
|---|---|---|---|---|
| Loreau and de Mazancourt (2008) | | 0 | 1 | 0.36 |
| Proposed | $q = 1$ | 0.992 | 0.998 | 0.985 |
| | $q = 2$ | 0.985 | 0.996 | 0.973 |

*Theoretical explanations for the above two interpretational problems*

The primary reason for the interpretational problems described above is that, for a measure to be interpretable, it must attain an extreme value (maximum or minimum) under a unique condition. As stated in the main text, our proposed measure of any order $q > 0$ attains its maximum or minimum under a unique condition, as stated in the main text:

- Our measures for any $q > 0$ attain a minimum value of 0 *if and only if* the biomass in the pooled dataset, at any time point, is almost exclusively contributed from one dataset, while the other datasets have negligible biomass.
- Our proposed synchrony measure for all $q > 0$ attains a maximum value of 1 *if and only if* all datasets have identical relative biomass vectors.

Thus, our synchrony measures clearly quantify the closeness/similarity between the multiple relative data vectors. However, the measure $\phi_{LM}$ attains its extremes under different conditions, as discussed earlier and detailed below.

- *The measure $\phi_{\mathrm{LM}}$ attains a maximum value of 1 under two different conditions*

Based on Eq. (S3.1), the measure $\phi_{\mathrm{LM}}$ attains its maximum value of 1 under two distinct conditions: (i) when the correlation between any two datasets is 1 (as illustrated in matrix $D_1$ below), or (ii) when only one dataset exhibits fluctuation, while all other datasets show no fluctuation (as illustrated in matrices $D_2$ and $D_3$ below). Since different conditions can lead to the same maximum value, $\phi_{\mathrm{LM}}$ does not provide an unambiguous quantification of the overall extent of pairwise correlation. Consequently, the intended target of quantification becomes unclear, leading to the two interpretational issues described above.

The two datasets in the following matrix $D_1$ exhibit a correlation of 1, implying $\phi_{\mathrm{LM}}$ = 1. Since the two datasets in matrix $D_1$ have the same relative vectors, our proposed synchrony measures ${}^1\psi$ and ${}^2\psi$ also attain the maximum value of 1. (Mathematically, it can be proved that when all datasets have identical relative biomass vectors, the correlation between any two datasets must be 1, but the reverse is not necessarily true.) Consider matrices $D_2$ and $D_3$, where only one dataset fluctuates. In both cases, $\phi_{\mathrm{LM}}$ attains the maximum value of 1 despite the fact that pairwise correlations in each matrix are not equal to 1 (and, in fact, are not well-defined, due to zero variance). Our proposed synchrony measures ${}^1\psi$ and ${}^2\psi$ are well-defined and yield values between 0 and 1, reflecting the differences in relative vectors within each matrix.

| Biomass matrix (row: dataset column: time) | | $D_1 = \begin{bmatrix} 1 & 2 & 5 & 8 \\ 2 & 4 & 10 & 16 \end{bmatrix}$ | $D_2 = A_1 = \begin{bmatrix} 5 & 5 & 5 & 5 \\ 5 & 5 & 5 & 4 \end{bmatrix}$ | $D_3 = B_1 = \begin{bmatrix} 2 & 2 & 2 & 2 \\ 3 & 3 & 3 & 3 \\ 1 & 4 & 5 & 8 \end{bmatrix}$ |
|---|---|---|---|---|
| Loreau and de Mazancourt (2008) | | 1 | 1 | 1 |
| Proposed | $q=1$ | 1 | 0.998 | 0.927 |
| | $q=2$ | 1 | 0.996 | 0.887 |

- *The measure $\phi_{\mathrm{LM}}$ attains a minimum value of 0 under multiple data structures*

The measure $\phi_{\mathrm{LM}}$ attains its minimum value of 0 when the total biomass in the pooled dataset remains constant over time. However, multiple data structures can fulfill this condition. For example, consider a matrix with two datasets, in which the pooled biomass remains constant at 10 over time; the following three data structures all meet this minimum condition:
(a) the two datasets are nearly identical, as illustrated in matrix $E_1$ below;
(b) the biomass in each dataset fluctuates, but at any time point, the biomass in the pooled dataset is contributed almost exclusively by one dataset, while the others contribute negligibly, as shown in matrix $E_2$;
(c) the biomass in each dataset fluctuates, and all datasets contribute to the pooled biomass at each time point, as shown in matrix $E_3$.

In all three data structures, $\phi_{\mathrm{LM}} = 0$, signifying that the measure fails to differentiate among the three structures. In contrast, for the nearly identical datasets in matrix $E_1$, our proposed synchrony measures ${}^1\psi$ and ${}^2\psi$ both approach 1, reflecting a high degree of synchrony (i.e., their relative vectors are very similar). The datasets in matrix $E_2$ satisfy the minimum condition for our measures, resulting in ${}^{1}\psi$ and ${}^{2}\psi$ both attaining the minimal value of 0. For matrix $E_3$, both ${}^{1}\psi$ and ${}^{2}\psi$ yield intermediate values between 0 and 1.

| Biomass matrix (row: dataset column: time) | | $E_1 = C_1 = \begin{bmatrix} 5 & 5 & 5 & 6 \\ 5 & 5 & 5 & 4 \end{bmatrix}$ | $E_2 = \begin{bmatrix} 0 & 0 & 10 & 10 \\ 10 & 10 & 0 & 0 \end{bmatrix}$ | $E_3 = \begin{bmatrix} 2 & 4 & 1 & 7 \\ 8 & 6 & 9 & 3 \end{bmatrix}$ |
|---|---|---|---|---|
| Loreau and de Mazancourt (2008) | | 0 | 0 | 0 |
| Proposed | $q = 1$ | 0.996 | 0 | 0.754 |
| | $q = 2$ | 0.985 | 0 | 0.589 |

*References cited in Appendix S3*


Loreau, M. and de Mazancourt, C. (2008). Species synchrony and its drivers: Neutral and nonneutral community dynamics in fluctuating environments. *American Naturalist*, 172, E48–E66.

Thibaut, L.M. and Connolly, S.R. (2013). Understanding diversity–stability relationships: Towards a unified model of portfolio effects. *Ecology Letters*, 16(2), 140–150.

Wang, S. and Loreau, M. (2014). Ecosystem stability in space: α, β and γ variability. *Ecology Letters*, 17, 891–901.

### Appendix S4. Data and Figures Referenced in the Applications Section

**Table S4.1**. The distribution of the number of species and plots in four blocks of the Jena Experiment (total number of plots = 76)

| Block | Block 1 | | | | | Block 2 | | | | | Block 3 | | | | | Block 4 | | | | |
|---|---|---|---|---|---|---|---|---|---|---|---|---|---|---|---|---|---|---|---|---|
| Number of species sown | 1 | 2 | 4 | 8 | 16 | 1 | 2 | 4 | 8 | 16 | 1 | 2 | 4 | 8 | 16 | 1 | 2 | 4 | 8 | 16 |
| Number of plots | 3 | 4 | 4 | 4 | 4 | 4 | 4 | 4 | 4 | 3 | 4 | 4 | 4 | 4 | 4 | 3 | 4 | 4 | 4 | 3 |
| Total number of plots | 19 | | | | | 19 | | | | | 20 | | | | | 18 | | | | |

**Table S4.2**. Biomass data from 2003 to 2024 for Plot B1A04 (4 species); species biomass data in 2004 were not available

| Year | 2003 | 2004 | 2005 | 2006 | 2007 | 2008 | 2009 | 2010 |
|---|---|---|---|---|---|---|---|---|
| Plot B1A04 | 726.75 | 1094.65 | 837.4 | 1092.575 | 1106.767 | 159.3533 | 426.0633 | 315.015 |
| Species Cam.pat | 4 | NA | 0.25 | 0 | 0 | 3.2 | 1.1667 | 0 |
| Species Fes.pra | 205.575 | NA | 472.4667 | 129.575 | 117.1675 | 75.0533 | 13.5633 | 20.515 |
| Species Ono.vic | 449.25 | NA | 314.375 | 800.075 | 762.8583 | 0 | 39 | 0 |
| Species Pla.lan | 67.925 | NA | 50.3083 | 162.925 | 226.7417 | 81.1 | 372.3333 | 294.5 |

| Year | 2011 | 2012 | 2013 | 2014 | 2015 | 2016 | 2017 | 2018 |
|---|---|---|---|---|---|---|---|---|
| Plot B1A04 | 172.425 | 172.295 | 246.845 | 233 | 248.465 | 213.75 | 313.28 | 112.18 |
| Species Cam.pat | 3.32 | 0 | 0 | 0 | 0 | 0 | 0 | 0 |
| Species Fes.pra | 49.105 | 49.580 | 176.295 | 182.45 | 145.95 | 189 | 6.4467 | 1.6433 |
| Species Ono.vic | 0 | 93 | 0 | 7.2 | 90.25 | 22.53 | 1.6 | 2.77 |
| Species Pla.lan | 120 | 29.715 | 70.55 | 43.35 | 12.265 | 2.22 | 305.2333 | 109.7667 |

| Year | 2019 | 2020 | 2021 | 2022 | 2023 | 2024 |
|---|---|---|---|---|---|---|
| Plot B1A04 | 633.66 | 269.17 | 379.775 | 291.685 | 368.5 | 577.39 |
| Species Cam.pat | 0 | 0 | 0 | 0 | 0 | 0 |
| Species Fes.pra | 20.305 | 0 | 0 | 46.5 | 0 | 0.98 |
| Species Ono.vic | 572.35 | 256.45 | 191.21 | 240 | 334.2 | 487.25 |
| Species Pla.lan | 41.005 | 12.72 | 188.565 | 5.185 | 34.3 | 89.16 |

**Table S4.3**. Biomass data from 2003 to 2024 for Plot B4A14 (2 species); species biomass data in 2004 were not available

| Year | 2003 | 2004 | 2005 | 2006 | 2007 | 2008 | 2009 | 2010 |
|---|---|---|---|---|---|---|---|---|
| Plot B4A14 | 192.6575 | 252.6 | 91.6908 | 144.7 | 134.08 | 133.1133 | 132.4167 | 201.265 |
| Species Bel.per | 1.2325 | NA | 3.4158 | 5.675 | 7.8383 | 11.1900 | 5.8833 | 13.765 |
| Species Pla.lan | 191.425 | NA | 88.2750 | 139.025 | 126.2417 | 121.9233 | 126.5333 | 187.5 |

| Year | 2011 | 2012 | 2013 | 2014 | 2015 | 2016 | 2017 | 2018 |
|---|---|---|---|---|---|---|---|---|
| Plot B4A14 | 224.26 | 71.765 | 53.8 | 73.875 | 62.51 | 4.875 | 125.2117 | 33.0467 |
| Species Bel.per | 5.76 | 12.765 | 23.95 | 3.175 | 14.755 | 2.005 | 0.975 | 0.1667 |
| Species Pla.lan | 218.5 | 59 | 29.85 | 70.7 | 47.755 | 2.87 | 124.2367 | 32.88 |

| Year | 2019 | 2020 | 202 | 2022 | 2023 | 2024 |
|---|---|---|---|---|---|---|
| Plot B4A14 | 5.56 | 14.7 | 68.455 | 82.4 | 53.87 | 82.505 |
| Species Bel.per | 0 | 0.49 | 4.305 | 4.05 | 1.07 | 5.44 |
| Species Pla.lan | 5.56 | 14.21 | 64.15 | 78.35 | 52.8 | 77.065 |

*Relationships between biodiversity and gamma stability, alpha stability, species synchrony and asynchrony across three time periods*

The three 10-year time periods considered here are as follows: an early period (2005–2014), a middle period (2010–2019), and a late period (2015–2024). The early and late periods are non-overlapping, while the middle period overlaps with each of the other two by five years. Figure S4.1 shows the relationships between biodiversity and gamma stability, alpha stability, species synchrony, and asynchrony across the three time periods.

- Panel (a) shows that the stabilizing effect of biodiversity on gamma stability strengthens over time, as indicated increasingly positive slopes.
- Panel (b) indicates that the destabilizing effect of biodiversity on alpha stability diminishes over time, because the magnitude of the negative slope diminishes and becomes positive in the late period, for $q = 0.5$ and $q = 1$.
- Panel (c) shows that the negative effect of biodiversity on species synchrony weakens over time, as indicated by the decreasing magnitude of the negative slope.
- Panel (d) reveals that the positive effect of biodiversity on species asynchrony also weakens over time, as indicated by progressively smaller positive slopes.

See the main text for general conclusions based on 12 consecutive 10-year moving windows.

(a) Relationship between biodiversity and gamma stability across three 10-year periods (positive slope decreases over time)

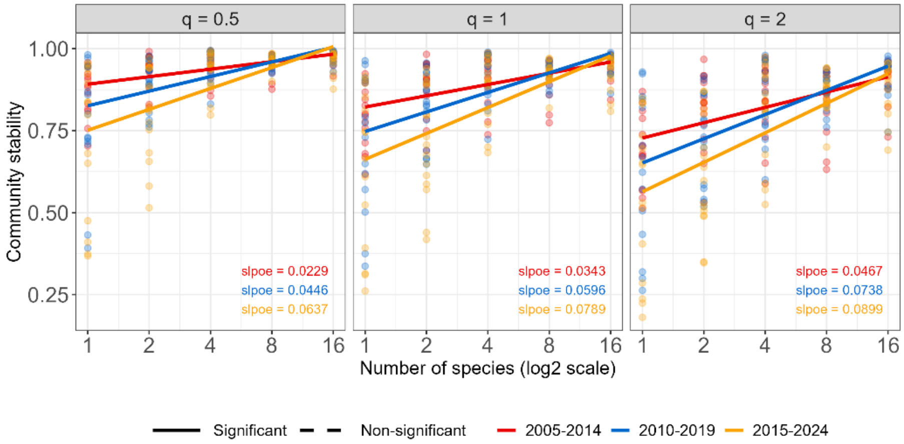


(b) Relationship between biodiversity and alpha stability across three 10-year periods (magnitude of negative slope diminishes and becomes positive in the late period for $q = 0.5$ and $q = 1$)

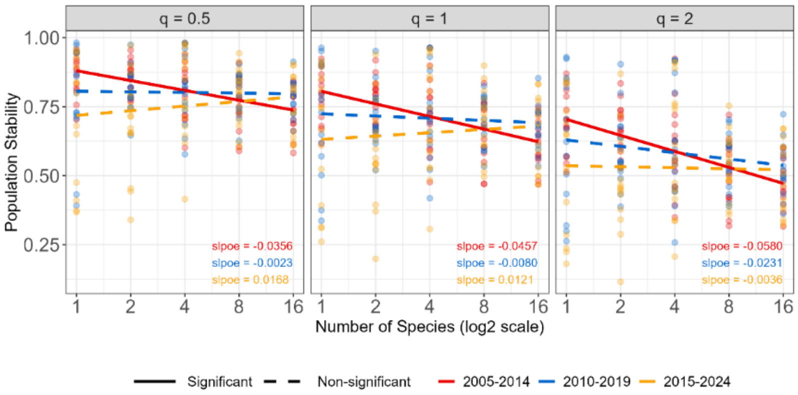


(c) Relationship between biodiversity and species synchrony across three 10-year periods (magnitude of negative slope decreases over time)

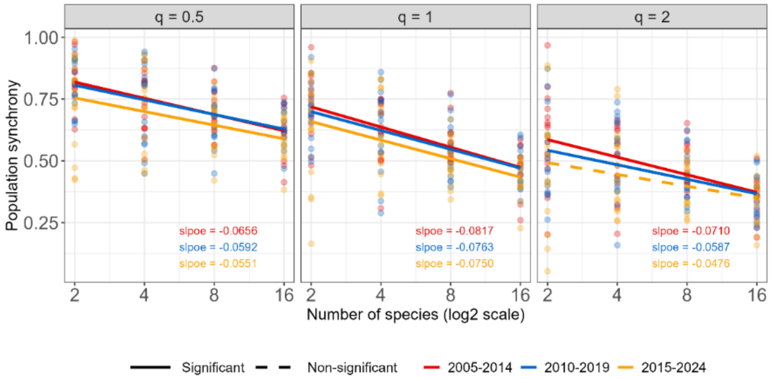


(d) Relationship between biodiversity and species asynchrony across three 10-year periods (positive slope decreases over time)

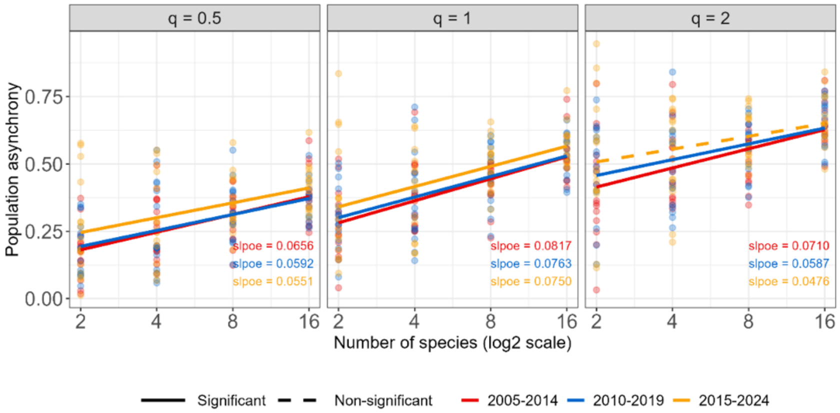


**Figure S4.1**. Relationships between biodiversity and (a) gamma stability, (b) alpha stability, (c) species synchrony, and (d) species asynchrony for three time periods: the early period (2005–2014), the middle period (2010–2019), and the late period (2015–2024).

#### Appendix S5. Proofs and Figures referenced in the Discussion Section

*The limiting behavior of some measures when time series length approaches infinity*

Assume that the biomass time series $\{z(1), z(2), ..., z(T)\}$ represent $T$ independent random variables drawn from a continuous non-negative random variable $X$ with probability density function (pdf) $f(x)$. Let $X$ have mean $\mu_f$, variance $\sigma_f^2$, coefficient of variation $CV_f^2 = \sigma_f^2 / (\mu_f)^2$ and entropy $H_f = E(-X \log X) = -\int (x \log x) f(x) dx$. The subscript $f$ indicates that these parameters describe properties of the underlying distribution, rather than statistics computed from the observed time series data. Following the notation in the main text, we denote $r(t) = z(t) / z$ as the relative biomass at time point $t$, where $z = z(+) = \sum_{t=1}^{T} z(t)$, and denote $\boldsymbol{r} = \{r(t); t = 1, 2, ..., T\}$ as the corresponding relative biomass vector.

**Theorem S5.1** As time series length $T$ approaches infinity, we can prove the following limiting behaviors:

(a) The two simple adjusted measures of stability, $H(\boldsymbol{r}) / \log T$ (for $q$ = 1) and $1 - CV^2(\boldsymbol{r}) / T$ (for $q$ = 2), both converge to a constant of 1.

(b) Our measure for $q$ = 1 converges to a limit expressed in terms of the mean and entropy of the underlying distribution. That is,

$$^1S(\boldsymbol{r}) = \frac{\exp[H(\boldsymbol{r})] - 1}{T - 1} \to \mu_f \exp\left(\frac{H_f}{\mu_f}\right).$$

(c) Our stability measure of $q$ = 2 converges to a limit expressed in terms of $CV_f^2$, i.e.,

$$^2S(\boldsymbol{r}) \to \frac{1}{CV_f^2 + 1} = \frac{\mu_f^2}{\mu_f^2 + \sigma_f^2}.$$

**Proof:**

(a) Based on Eq. (2.1c) of the main text, we can rewrite $H(\boldsymbol{r}) / \log T$ as

$$\frac{H(\boldsymbol{r})}{\log T} = -\frac{1}{\log T} \sum_{t=1}^{T} r(t) \log r(t) = -\frac{1}{\log T} \sum_{t=1}^{T} \frac{z(t)}{z} \log\left(\frac{z(t)}{z}\right)$$

$$= \frac{1}{\log T}\left[-\sum_{t=1}^{T} \frac{z(t) \log z(t)}{z} + \log \frac{z}{T} + \log T\right]$$

$$= 1 + \frac{1}{\log T}\left[-\sum_{t=1}^{T} \frac{z(t) \log z(t)}{z} + \log \frac{z}{T}\right].$$

The law of large numbers implies that the two terms in the brackets of the above formula converge to finite values because $\log \frac{z}{T} \to \log \mu_f$ and

$$-\sum_{t=1}^{T} \frac{z(t)\log z(t)}{z} = -\sum_{t=1}^{T} \frac{z(t)\log z(t)/T}{z/T} \to \frac{H_f}{\mu_f}.$$

This result implies that $H(\boldsymbol{r})/\log T$ converges to 1. For $q = 2$, it follows again from the law of large numbers that $CV^2(\boldsymbol{r}) \to CV_f^2$, implying that $1 - CV^2(\boldsymbol{r})/T$ also tends to 1.

(b) Note that we have the following expression for $\exp[H(\boldsymbol{r})]$:

$$\begin{aligned}\exp[H(\boldsymbol{r})] &= \exp\left[-\sum_{t=1}^{T} \frac{z(t)}{z} \log\left(\frac{z(t)}{z}\right)\right] \\ &= \exp\left[-\sum_{t=1}^{T} \frac{z(t)\log z(t)}{z} + \sum_{t=1}^{T} \frac{z(t)}{z}\log z\right] \\ &= z \times \exp\left[-\sum_{t=1}^{T} \frac{z(t)\log z(t)}{z}\right].\end{aligned}$$

By the law of large numbers, we obtain

$$\frac{\exp[H(\boldsymbol{r})]}{T} = \frac{z}{T} \times \exp\left[-\sum_{t=1}^{T} \frac{z(t)\log z(t)}{z}\right] \to \mu_f \exp\left(\frac{H_f}{\mu_f}\right).$$

Then

$${}^1S(\boldsymbol{r}) = \frac{\exp[H(\boldsymbol{r})] - 1}{T - 1} \sim \frac{\exp[H(\boldsymbol{r})]}{T} \to \mu_f \exp\left(\frac{H_f}{\mu_f}\right).$$

(c) As stated in (a), we have $CV^2(\boldsymbol{r}) \to CV_f^2$. The limit, below, then follows from Eq. (2.3b) of the main text.

$${}^2S(\boldsymbol{r}) = \frac{1 - \frac{CV^2(\boldsymbol{r})}{T}}{CV^2(\boldsymbol{r}) + \left(1 - \frac{CV^2(\boldsymbol{r})}{T}\right)} \to \frac{1}{CV_f^2 + 1} = \frac{\mu_f^2}{\mu_f^2 + \sigma_f^2}. \text{ QED}$$

**Theorem S5.2.** In the special case that $X$ follows a gamma distribution with pdf

$$f(x) = x^{k-1} e^{-x/\theta} / [\Gamma(k)\theta^k], \quad k, \theta > 0.$$

Then ${}^2S(\boldsymbol{r}) \to \frac{1}{(1/k)+1}$, and ${}^1S(\boldsymbol{r}) \to k\exp[-\psi(k+1)]$, where $\psi$ denotes the digamma function.

**Proof:** Under the gamma distribution, we have mean $\mu_f = k\,\theta$, variance $\sigma_f^2 = k\theta^2$, and $CV_f^2 = \sigma_f^2/(\mu_f)^2 = 1/k$. Thus, ${}^2S(\mathbf{r}) \to \dfrac{1}{(1/k)+1}$ follows from part (c) of Theorem S5.1. For the proposed measure for $q$ = 1, note that

$$H_f = -\int (x \log x) x^{k-1} e^{-x/\theta} / [\Gamma(k)\theta^k] dx$$

$$= -k\theta \int (\log x) x^k e^{-x/\theta} / [\Gamma(k+1)\theta^{k+1}] dx = -k\theta \times E(\log Y),$$

where $Y$ follows a gamma distribution with parameter $k$+1 and $\theta$. A direct derivation yields the following result: $E(\log Y) = \psi(k+1) + \log\theta$. Then we obtain, from (b) of Theorem S5.1, that

$${}^1S(\mathbf{r}) = \frac{\exp[H(\mathbf{r})]-1}{T-1} \to \mu_f \exp\left(\frac{H_f}{\mu_f}\right) = k\exp[-\psi(k+1)]. \text{ QED}$$

*Plot within-species population stability with respect to species weight*

**(a)**

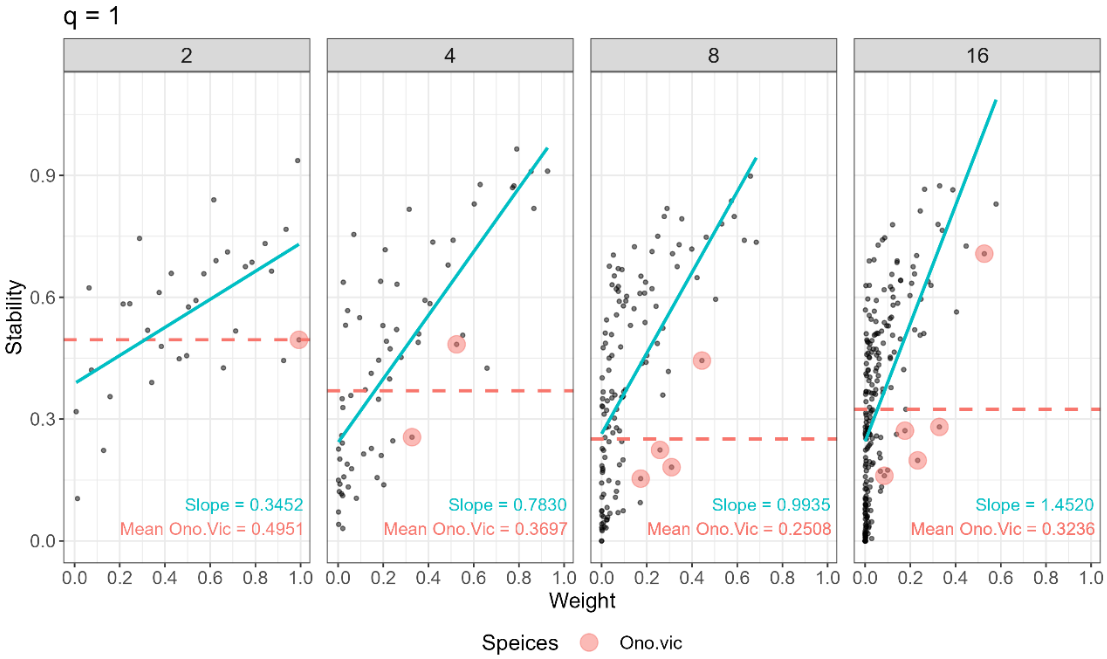

**(b)**

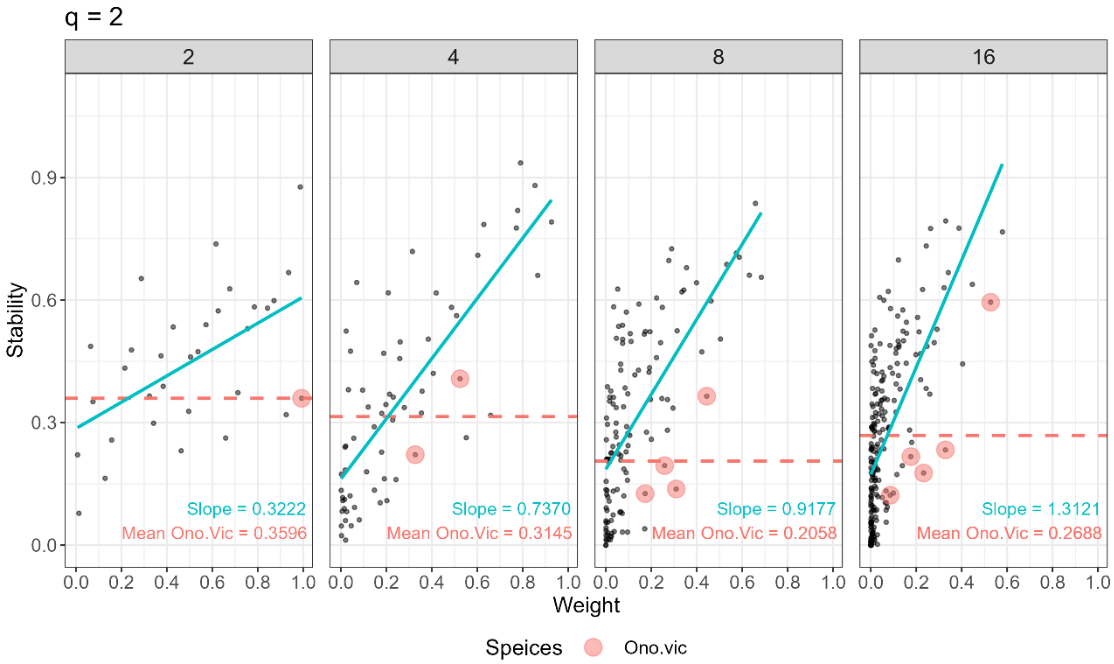


**Figure S5.** The linear relationships between within-species population stability and species relative-biomass weight in a plot for (a) $q$ = 1, (b) $q$ = 2, for species = 2, 4, 8, and 16 (Column 1 to Column 4). Pink dots represent the stability values for a dominant species (*Onobrychis viciifolia Scop.*, which belongs to the functional class of legumes). The average values of stability for this dominant species are shown.
